# A proprioceptive feedback circuit drives *C. elegans* locomotor adaptation through dopamine signaling

**DOI:** 10.1101/2022.10.14.512295

**Authors:** Hongfei Ji, Anthony D. Fouad, Zihao Li, Andrew Ruba, Christopher Fang-Yen

## Abstract

An animal adapts its motor behavior to navigate the external environment. This adaptation depends on proprioception, which provides feedback on an animal’s body postures. How proprioception mechanisms interact with motor circuits and contribute to locomotor adaptation remains unclear. Here we describe and characterize proprioception-mediated homeostatic control of undulatory movement in the roundworm *Caenorhabditis elegans*. We found the worm responds to optogenetically or mechanically induced decreases in midbody bending amplitude by increasing its anterior amplitude. Conversely, it responds to increased midbody amplitude by decreasing the anterior amplitude. Using genetics, microfluidic and optogenetic perturbation response analyses, and optical neurophysiology, we elucidated the neural circuit underlying this compensatory postural response. The dopaminergic PDE neurons proprioceptively sense midbody bending and signal to AVK interneurons via the D2-like dopamine receptor DOP-3. The FMRFamide-like neuropeptide FLP-1, released by AVK, regulates SMB head motor neurons to modulate anterior bending. We propose that this homeostatic behavioral control optimizes locomotor efficiency. Our findings demonstrate a mechanism in which proprioception works with dopamine and neuropeptide signaling to mediate motor control, a motif that may be conserved in other animals.

## Introduction

Animal navigation in complex natural environments requires flexible locomotor behavior (1). During locomotion, an animal needs to adapt its body posture and motor output to its surrounding context and the inevitable perturbations resulting from obstacles and irregularities (2). Kinematic and electromyographic studies in legged animals revealed phasic compensatory reactions in their central nervous systems, characterized by rapid corrective movements adapted to the perturbation (3, 4). In undulatory animals such as fish and nematodes, motor circuits also contextually tune motor behaviors to external conditions (5–7). The observations of these adaptive movements in various conditions highlight the behavioral flexibility of motor systems across species.

In many animals, adaptive locomotory movements involve interactions between neural circuits called central pattern generators (CPGs) capable of generating primary locomotor rhythms (8–11) and sensory feedback that modulates locomotion (2, 12–14). In particular, proprioception provides rapid feedback on body position for locomotor control during movements (15–17). In mammals, proprioceptive inputs from multiple sensory organs are continuously processed within spinal cord circuits to adapt motor behavior (18). In lamprey and other undulatory animals, locomotion also relies on proprioceptive feedback to adjust the locomotor pattern to changes in the physical environment (19–21).

The mechanisms by which the nervous system controls adaptive locomotion are complex and poorly understood. In vertebrates, corrective locomotor control involves multiple circuits in the spinal cord, brainstem, and forebrain that process sensory feedback and generate appropriate motor commands (22–25). Recent advances in genetic techniques have improved the ability to study the function of CPGs and their constituent neurons in fine motor control (12, 26). For example, in mice, many spinal interneurons have been identified as critical components of the corrective locomotor control system (27). However, it is still unclear how these neurons contribute to locomotor movements, partly due to the lack of *in vivo* methods for acutely perturbing their activity (2). Furthermore, our understanding of how proprioceptive input is integrated and transmitted to control movement and posture and which circuit components are responsible for relaying these signals remains limited (28).

Here, we use the roundworm *C. elegans* to examine how corrective locomotor control is executed by the nervous system. *C. elegans* has a relatively small and well-characterized nervous system with fully identified cell types and mapped synaptic connectivity (29, 30). A wealth of methods for assaying and manipulating *C. elegans* (31–33) offers the unique opportunity for an integrative dissection of locomotor control at the systems, circuit, cellular, and molecular genetic levels.

*C. elegans* moves forward by propagating dorso-ventral bending waves from anterior to posterior. These undulatory movements are generated by body wall muscles arranged in dorsal and ventral rows along the worm’s body (34). The alternating activity of antagonistic muscles is driven by motor neurons located in the head ganglia and the ventral nerve cord (35). A set of premotor interneurons is responsible for coordinating forward and reverse locomotion (36).

While motor neurons and premotor interneurons generate the worm’s sinusoidal bending (30, 37), adaptive locomotion in natural contexts requires motor control of the head and involves sublateral motor neurons that modulate posture (38–40). Optimal motor control is context dependent and subject to feedback input from a large number of interneurons and sensory neurons (41–45). Biogenic amine and neuropeptide neuromodulators are also involved in various long-term and short-term locomotor states (46–51).

Proprioception plays a critical role in the motor behavior of *C. elegans*. Several classes of neurons have been identified as proprioceptors important for the worm’s locomotor patterns. The B-type motor neurons facilitate proprioceptive coupling from anterior to posterior bending to propagate undulatory waves along the body (52). The head motor neurons SMDD regulate head steering movement during locomotion (40) and have been proposed as proprioceptors and candidate locomotor CPG elements (39). In addition, the DVA and PVD interneurons are also involved in regulating the worm’s body bend movement through proprioceptive mechanisms. DVA acts as a proprioceptor by relying on the mechanosensitive channel TRP-4 to regulate the body bend amplitude (43). PVD releases the NLP-12 neuropeptide from its dendrites when it is proprioceptively triggered by local body bending, which is thought to regulate the amplitude of body movements (45).

In this study we used physical and optical perturbations to further investigate how the *C. elegans* motor system controls its bending amplitude during locomotion. Using targeted optogenetic manipulation in freely moving animals, we found that *C. elegans* uses a posterior-to-anterior proprioceptive feedback loop to adapt its locomotor amplitude to perturbations. We dissected the underlying neuronal pathway, showing that dopaminergic PDE neurons proprioceptively respond to midbody curvature and drive AVK interneuron activity via the D2-like dopamine receptor DOP-3. The neuron AVK, in turn, releases the FLP-1 FMRFamide-like neuropeptide, which regulates downstream SMB head motor neurons to modulate anterior bending. Our findings reveal a circuit for adaptive movement control by *C. elegans* from sensory input to motor output. We discuss how this feedback control mechanism might help optimize locomotor efficiency and foraging behavior.

## Results

*C. elegans* bidirectionally modulates anterior bending amplitude in response to optogenetic perturbation of midbody curvature.

To measure locomotor behavior, we calculate curvature along the worm body centerline (6) (Fig. 1A). With this metric we quantified undulatory movements of freely moving *C. elegans* via the time-varying normalized curvature from head to tail (Fig. 1C, F).

**Figure 1.**
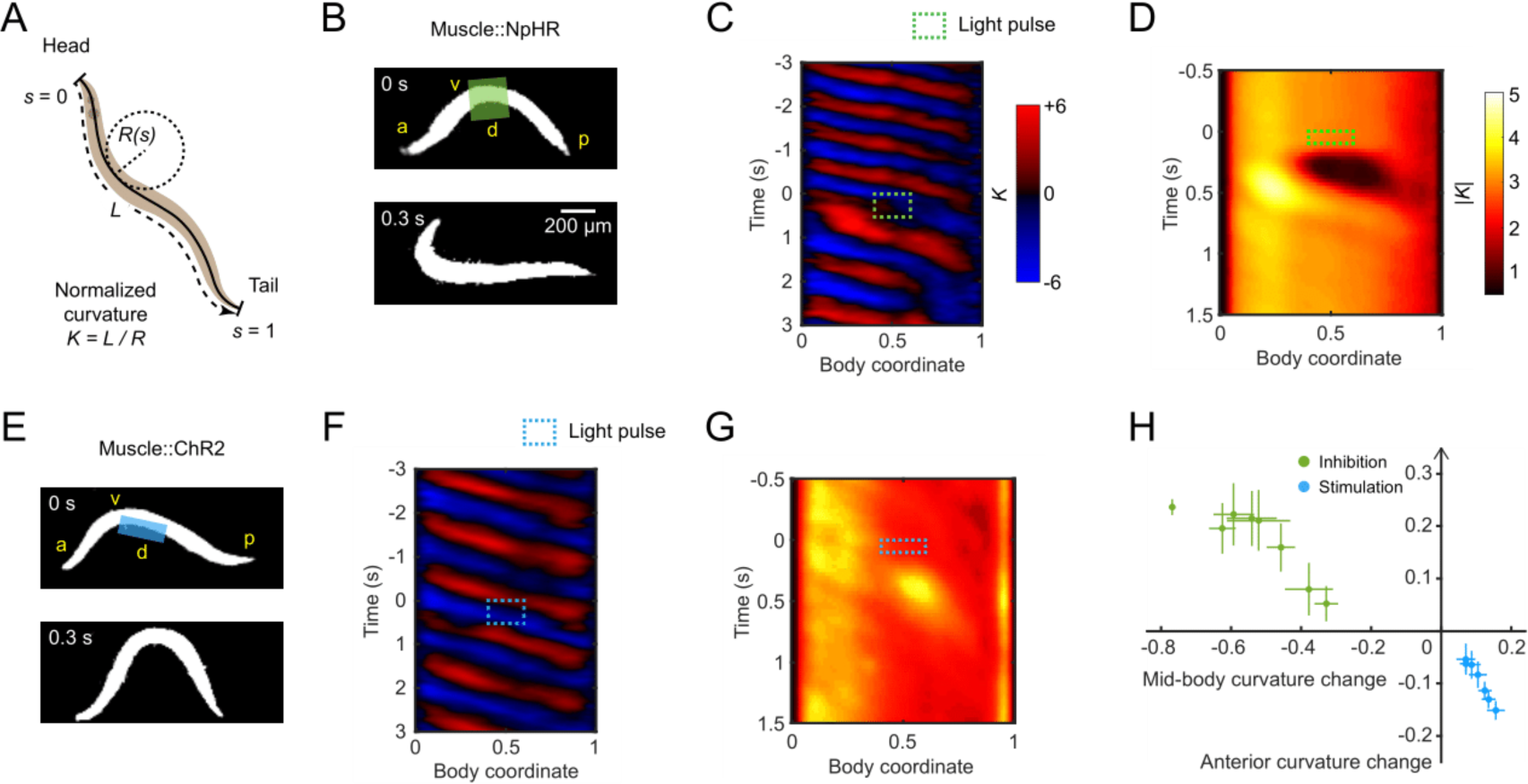
*C. elegans* bidirectionally modulates anterior bending amplitude in response to optogenetic perturbation of midbody curvature. (A) Quantification of worm locomotion using time-varying curvature. Body coordinate 𝑠 is denoted by the distance along the centerline normalized by the body length *L* (head = 0, tail = 1). Normalized curvature 𝐾 is the product of the body length and the reciprocal local radius of body curvature 𝑅(𝑠), with positive and negative values representing ventral and dorsal bending, respectively. (B) Optogenetic muscle inhibition in the midbody during forward locomotion. Green region indicates the laser illumination. a: anterior, p: posterior, d: dorsal, v: ventral. (C) Curvature of the worm locomotion shown in (B). Green box indicates a 0.5 s laser illumination starting at 𝑡 = 0 applied to the midbody. (D) Mean absolute curvature around 0.1 s inhibitions (green box). n = 1160 midbody illuminations from 206 worms. (E) Optogenetic muscle stimulation in the dorsal midbody during forward locomotion. Blue region indicates the laser illumination. (F) Curvature of the worm locomotion shown in (E). Blue box indicates a 0.5 s laser illumination starting at 𝑡 = 0 applied to the dorsal midbody. (G) Mean absolute curvature around 0.1 s stimulations (blue box). n = 693 dorsal midbody illuminations from 122 worms. (H) Relationship between the mean normalized curvature change in the anterior vs. midbody regions. Each point represents mean ± SEM of the corresponding normalized value of the first post-illumination curvature peak. Green and blue points denote data induced by optogenetic midbody muscle inhibition (both sides) and stimulation (dorsal or ventral side), respectively. n = 110-150 illuminations from 10-20 animals per group.

In previous studies we perturbed the muscular or neural activity in different body regions and analyzed the resulting undulatory dynamics during forward locomotion (19). Using a laser targeting system (33, 53) we applied green laser illumination (532 nm wavelength) to selected body regions of animals expressing the inhibitory opsin *NpHR* in body wall muscles (via the *Pmyo-3* promoter).

Transient optogenetic inhibition (0.1 or 0.5 s duration) of muscles at the head region (0.05-0.25 body length) or neck region (0.2-0.4 body length) both caused rapid straightening of the anterior region followed by a mild amplitude decline in the subsequent body bends propagating from head to tail (Fig. S1A, B, E, F; Movie 1), consistent with previous findings (19).

When we inhibited muscles at the midbody (0.4-0.6 body length), in addition to observing a paralytic effect propagating from midbody to tail, we found worms exhibited exaggerated undulations at the anterior region (0.1-0.3 body length; Fig. 1C, D and S1G; Movie 1). Worms returned to the baseline undulatory amplitude within about one undulatory cycle after the inhibition (Fig. S1I, J).

We hypothesized that the increase in head bending amplitude represents a homeostatic response to the decrease in midbody curvature. To test this idea, we asked how the animal would react to an optogenetically-induced increase in midbody amplitude. We stimulated midbody muscles on the dorsal or ventral side by exposing the corresponding region of animals expressing the excitatory opsin *ChR2* in the body wall muscles to blue laser illumination (473 nm wavelength). Stimulating one side of the midbody muscles led to exaggerated midbody bending but a decreased anterior bending response (Fig. 1E-G and S1K, L; Movie 2). These data show that changes in anterior amplitude occur in the opposite direction to changes in midbody amplitude.

We also tested worm locomotion perturbed by brief muscle inhibition (0.1 s duration) at the posterior region (0.6-0.8 body coordinate). Posterior bending amplitude rapidly decreased upon illumination, but the bending amplitude of the anterior half did not increase (Fig. S1D, H; Movie 1). This result suggests that the anterior body bending responds to perturbations in the midbody but not the posterior region.

We compared the sensitivity of the anterior bending curvature in response to stimulation versus inhibition in the midbody muscles. We applied laser pulses to the midbody with varied pulse duration and irradiance to change the degree of muscle stimulation or inhibition (see *Materials and Methods*). With varying light dosage, changes in the anterior amplitude varied continuously in response to the induced changes in the midbody amplitude (Fig. 1H).

These observations indicate a mechanism that mediates anterior bending curvature in response to changes in midbody curvature. Because the anterior curvature can be thought of as compensating for changes in midbody curvature, we will henceforth refer to this effect as compensatory curvature response (CCR).

### Physical constraint of midbody causes an increase in anterior bending amplitude

We hypothesized that CCR requires proprioceptive sensing of midbody curvature. In our optogenetic experiments, curvature change was induced by muscle activity manipulation, leaving open the possibility that CCR arises from non-proprioceptive signaling within muscles or from muscles to neurons.

To test the involvement of proprioception in CCR, we designed a straight-channel microfluidic device that mechanically reduces midbody bending amplitude by partially constraining this region (Fig. 2A and S2A). In this manner, we manipulated body curvature without directly perturbing muscle activity and compared the perturbed behavior with normal locomotion (Fig. 2B).

**Figure 2.**
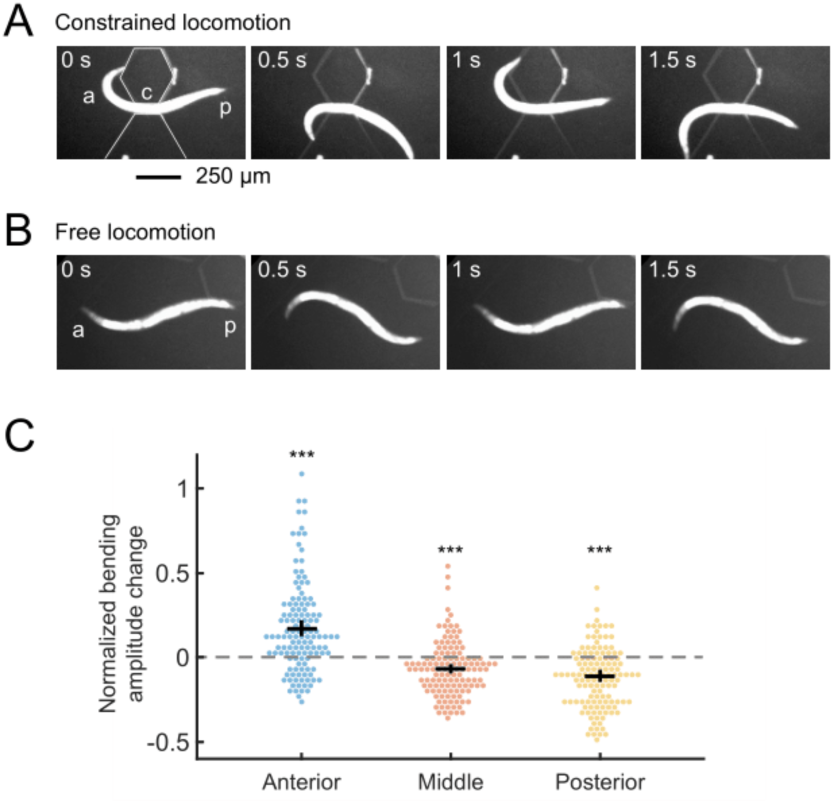
Physical constraint of midbody causes an increase in anterior bending amplitude. (A) A wild-type animal showing constrained locomotion with its midbody confined by a straight-channel microfluidic device. a: anterior region, p: posterior region, c: constrained middle region of a worm. (B) A wild-type animal during free locomotion in the wide region of the channel. (C) Effects of midbody constraint during forward locomotion on the undulatory bending amplitude of the anterior, middle, and posterior regions, measured as the normalized curvature change of the corresponding regions. n = 19 wild-type animals, mean ± SEM. ***p < 0.001 compared with zero curvature change (gray line).

During physical constraint, animals exhibited reduced bending amplitude in the constrained middle region and increased amplitude in the anterior region (Fig. 2C and S2B; Movie 3), which is comparable to the bending amplitude changes observed during optogenetic muscle inhibition of the midbody. These results suggest that CCR is reliant on proprioception.

In *C. elegans*, electrical synapses composed of the innexin UNC-9 connect adjacent body wall muscle cells (54). To examine whether these connections are involved in CCR, we assessed the role of UNC-9 in this process. UNC-9 is expressed in both the body wall muscles and the nervous system. Using the microfluidic device, we assayed transgenic animals that lacked functional UNC-9 in muscle cells but had normal UNC-9 expression in the nervous system (54). These animals exhibited normal CCR (Fig. S3B), suggesting that direct inter-muscular coupling is not required for CCR.

### Compensatory curvature response requires functional dopamine signaling by PDE neurons

To understand the mechanisms underlying CCR, we conducted a candidate gene screen using the straight-channel microfluidic device. For each candidate gene we evaluated the anterior bending amplitude of corresponding mutants constrained by the straight channel in the midbody and compared it with the amplitude during free locomotion. To quantify the extent of CCR, we computed a CCR index equal to the difference between the anterior bending amplitudes during constrained and free locomotion divided by the amplitude during free locomotion (see *Materials and Methods* for details).

To determine which neurotransmitter systems might be required for CCR, we analyzed mutant strains with deficiencies in the biosynthesis of biogenic amines, including dopamine, serotonin, tyramine, octopamine, and gamma-aminobutyric acid (GABA).

Mutants for *tph-1(n4622)* (defective in serotonin synthesis) and *tdc-1(n3421)* (defective in both tyramine and octopamine syntheses) displayed largely intact CCR during midbody constraint (Fig. S3B), suggesting that serotonin, tyramine, and octopamine are not required for CCR. Mutants *unc-25(e156)* (defective in GABA synthesis) displayed uncoordinated loopy locomotion when moving freely and a compromised CCR (Fig. S3B). By contrast, dopamine-deficient *cat-2(e1112)* mutants showed normal locomotion but greatly impaired CCR (Fig. 3A-C). The deficiency in *cat-2* CCR was fully restored by the addition of exogenous dopamine (Fig. 3C), implying that the defect in CCR in *cat-2* mutants is due to the lack of dopamine.

**Figure 3.**
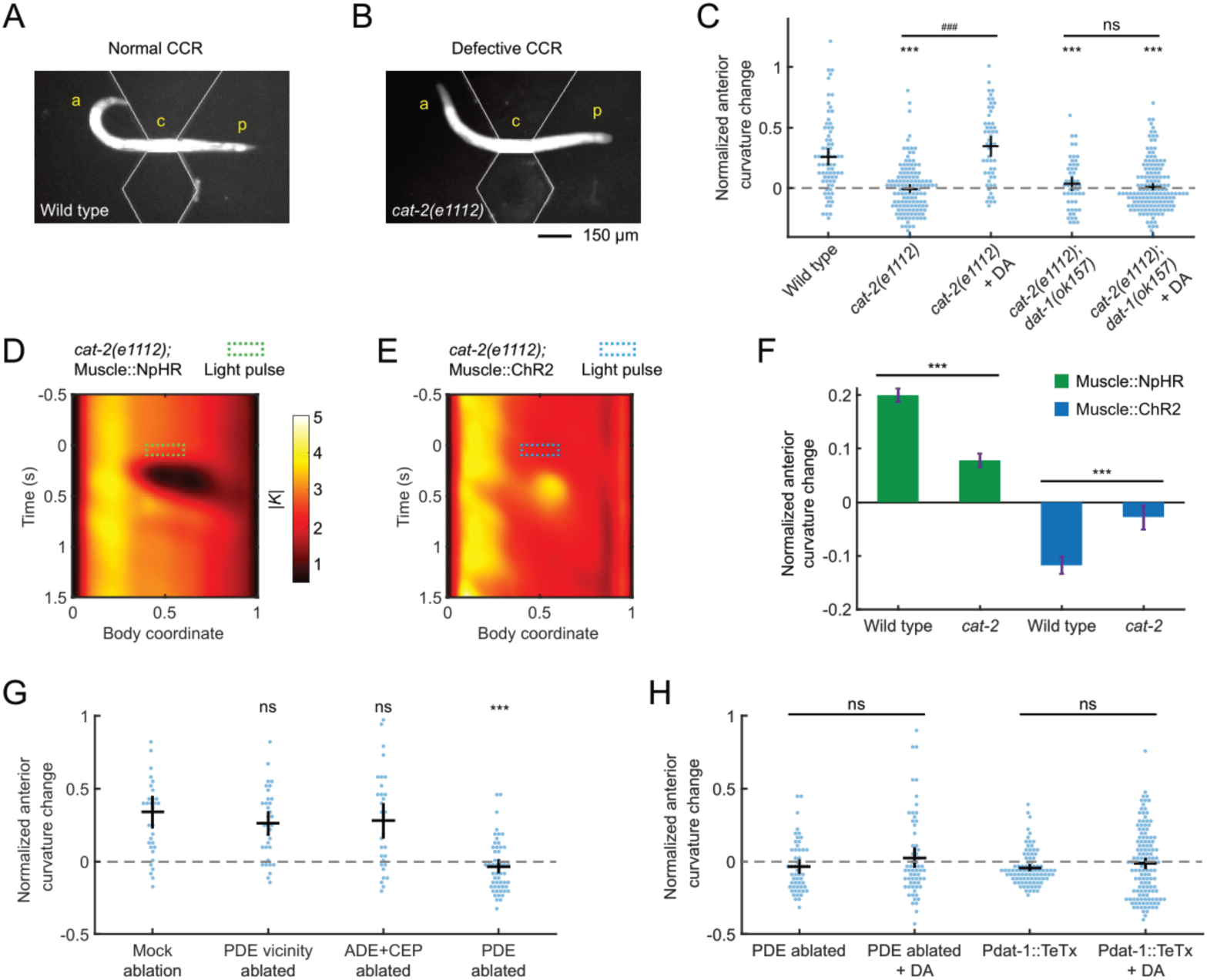
Compensatory curvature response requires functional dopamine signaling by PDE neurons. (A, B) A wild-type animal (A) and a *cat-2(e1112)* (B) mutant with midbody confined by a straight-channel microfluidic device, exhibiting undulations with normal and defective CCR, respectively. a: anterior region, p: posterior region, c: constrained middle region of a worm. (C) CCR indices for wild-type animals, mutant *cat-2(e1112)* and double mutant *cat-2*(*1112*)*; dat-1(ok157)* in either the absence or presence of exogenous 50 mM dopamine. n ≥ 10 animals per group. Error bars show mean ± SEM. ***p < 0.001 compared with wild type; ^###^p < 0.001 compared with *cat-2* mutants without exogenous dopamine; ns: not significant, compared with *cat-2; dop-3* mutants without exogenous dopamine, Tukey-Kramer multiple comparison tests. (D, E) Mean absolute curvature around 0.1 s illuminations (dashed box) for *cat-2* mutants expressing Muscle::NpHR (n = 133 midbody illuminations from 33 worms) or Muscle::ChR2 (n = 112 dorsal midbody illuminations from 24 worms). (F) Normalized anterior curvature change of the first post-illumination curvature peak for wild-type and *cat-2* animals expressing Muscle::NpHR (green) or Muscle::ChR2 (blue), mean ± SEM. ***p < 0.001, Student’s t-test. (G) CCR indices for animals with genetic ablation of ADEs and CEPs (Pdat-1::ICE, PDE survival confirmed by co-expression of Pdat-1::RFP), laser ablation of PDE vicinity regions, and laser ablation of PDEs, compared with a mock-ablation group. n ≥ 10 animals per condition, mean ± SEM. ***p < 0.001, ns: not significant, Dunnett’s multiple comparison tests. (H) CCR indices for PDE-ablated worms and transgenic animals expressing tetanus toxin light chain in all dopaminergic neurons (Pdat-1::TeTx), in the absence and presence of exogenous 50 mM dopamine, respectively. n ≥ 10 animals per condition. Error bars show mean ± SEM. ns: not significant, Student’s t-test.

In the experiments described above, the worm’s midbody curvature change was induced by either optogenetic manipulation or microfluidic constraint. While both procedures modify the animal’s midbody curvature, it is possible that the resulting changes in anterior curvature are due to distinct mechanisms. To investigate whether the mechanism for CCR is the same in response to these two manipulations, we induced midbody curvature changes using optogenetic manipulations in *cat-2* mutant animals that are defective in CCR in the microfluidic device. We crossed the transgenic animals expressing Muscle::NpHR or Muscle::ChR2 into a *cat-2* mutant background and performed the optogenetic muscle perturbation experiments as described above. Mutant animals *cat-2* exhibited impaired CCR to midbody curvature manipulation by optogenetic muscle inhibition or stimulation (Fig. 3D-F), indicating that dopamine signaling is required for CCR in response to both midbody curvature decrease and increase. These findings suggest that CCR reflects the same mechanism as observed in the microfluidic device and optogenetic experiments.

Since dopaminergic neuromodulation has been shown to play a role in *C. elegans* locomotion under food-related conditions (50), we examined whether CCR depends on food presence. Using the microfluidic device we compared CCR of wild-type animals under no food or two concentrations of food bacteria. We found that CCR did not depend on the amount of food (Fig. S2C).

Dopamine regulates a variety of behaviors in *C. elegans*, including locomotion (55), touch sensation (56), egg-laying (41), and gait transitions (51). The *C. elegans* hermaphrodite has 8 dopaminergic neurons consisting of 4 CEPs, 2 ADEs, and 2 PDEs (57). To investigate which neurons are required for CCR, we ablated specific subsets of dopaminergic neurons in L3 larvae and examined their responses to physical midbody constraints at the adult stage.

We first ablated ADEs and CEPs using transgenic animals expressing the human caspase interleukin-1β-converting enzyme (ICE) in dopaminergic neurons under the *dat-1* promoter (48). After crossing the *Pdat-1::ICE* strain with a transgene expressing mCherry in all dopaminergic neurons, we confirmed the cell death of ADEs and CEPs and noted the survival of PDEs by imaging the resulting RFP expression (see *Materials and Methods*). Next, we ablated only PDE somas or the vicinity region of PDEs using a focused infrared laser beam (58) (see *Materials and Methods*). Transgenic worms with ADEs and CEPs killed or PDE vicinity region damaged exhibited normal CCR, while worms lacking only PDEs were defective in CCR, and CCR could not be restored by exogenous dopamine (Fig. 3G, H). These results show that PDEs are the only dopaminergic neurons necessary for CCR.

Next we investigated double mutant animals *cat-2(e1112);dop-3(ok157)*, deficient in both dopamine biosynthesis and reuptake, and transgenic animals with dopaminergic neurons expressing tetanus toxin light chain (*Pdat-1::TeTx*), which blocks synaptic transmission (59). We observed compromised CCR in both strains, and supplementation with exogenous dopamine failed to restore their mutant phenotypes (Fig. 3C, H). Our findings suggest that local, timely release of dopamine from PDE is necessary for CCR, as restoration of CCR was only observed in dopamine synthesis-deficient *cat-2* mutants in exogenous dopamine environments but not in animals with deficits in dopamine reuptake, PDE elimination, or TeTx-expressing dopaminergic neurons.

To further investigate the role of PDE in CCR, we conducted optogenetic manipulations of PDE in transgenic animals expressing the excitatory opsin *GtACR2* or the inhibitory opsin *CoChR* (41) in all dopaminergic neurons (via the *Pdat-1* promoter). To target only PDE and avoid ADE and CEP, which are located in the head, we illuminated transgenic animals in the posterior region (0.4-0.8 body region). Optogenetically inhibiting or stimulating PDE in freely behaving animals resulted in increases or decreases in anterior curvature, respectively (Fig. S4).

Taken together, our findings suggest that functional dopamine release from PDE, but not from ADE or CEP, is necessary for CCR. The increase or decrease of anterior curvature amplitude during CCR may be linked to the deactivation or activation of PDE neurons, respectively.

### The dopaminergic PDE neurons respond to midbody curvature

The PDE neuron pair is the sole dopaminergic neuron type in *C. elegans* with neurites that extend across the midbody (30), which is the area where the curvature perturbation was applied. This fact and our discovery that PDE neurons are required for CCR prompted us to ask whether PDEs might function as proprioceptors for midbody curvature. A previous study revealed that PDE calcium activity in wild-type animals is synchronized with their bending waves during roaming (41); however, whether PDEs respond to body movements or function as proprioceptors remain unknown.

As a first step toward establishing a proprioceptive function for PDE, we monitored the spontaneous calcium transients of PDEs in freely crawling animals expressing the genetically encoded calcium indicators GCaMP in PDEs via the *Pdat-1* promoter (41) (Fig. 4A; see *Materials and Methods*). We observed robust oscillating calcium dynamics in the PDE somas during *C. elegans* forward movement (Fig. 4B). We also noticed that the calcium activity in PDE somas was correlated with the animal’s body curvature (Fig. 4E). These correlations between PDE fluorescence and body posture were not observed in transgenic animals expressing GFP in PDEs (Fig. S5). Our calcium imaging experiments indicate that the native neuronal activity of PDEs correlates with body posture during free locomotion of an intact wild-type animal, as previously reported (41).

**Figure 4.**
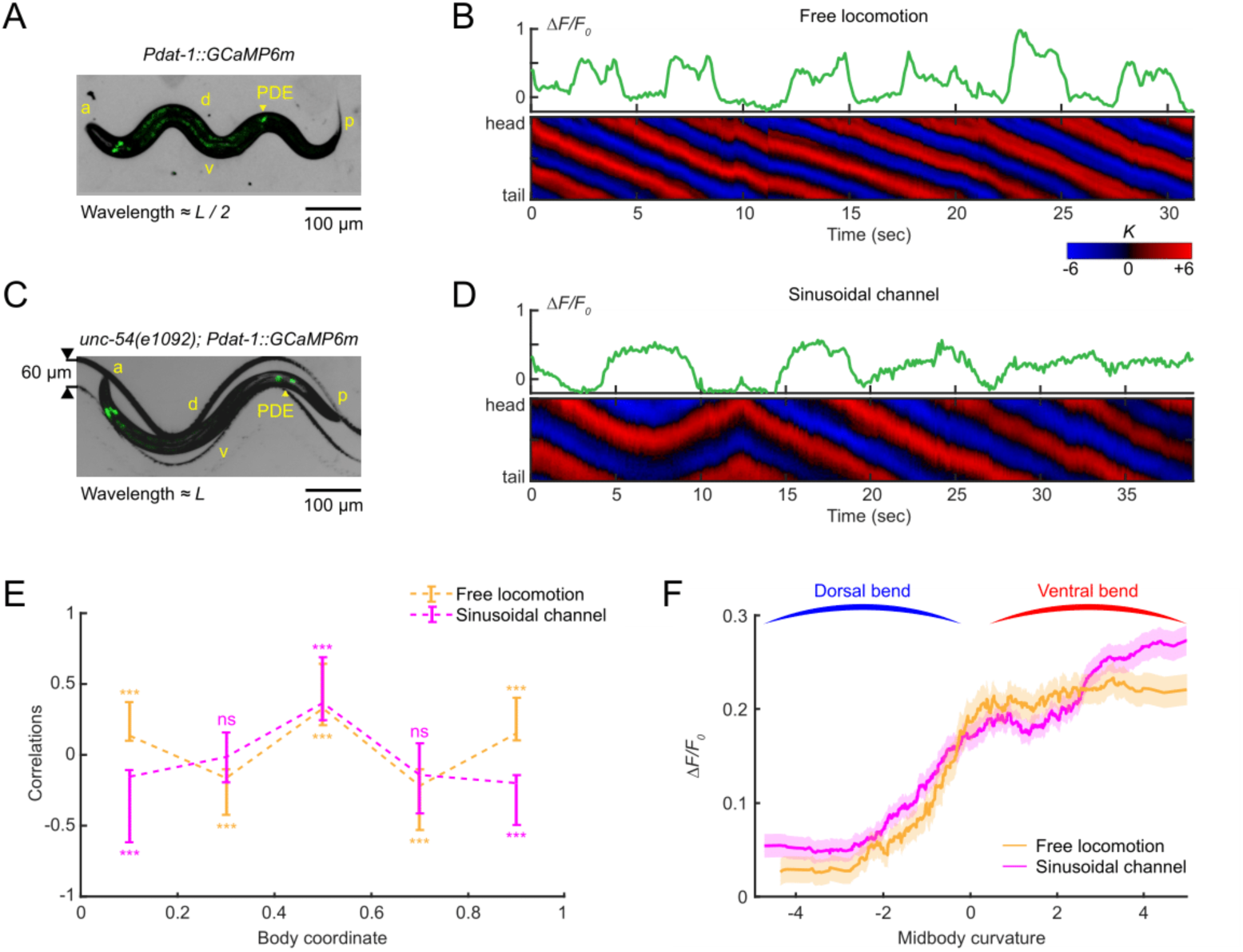
The dopaminergic PDE neurons respond to midbody curvature. (A, C) Overlay of a bright field and fluorescence image of the PDE neurons (via transgenic expression Pdat-1::GcaMP6m) in a freely moving wild-type animal on an agar surface (A) and a muscularly paralyzed mutant *unc-54(e1092)* within a 60-µm-wide sinusoidal channel (C). a: anterior, p: posterior, d: dorsal, v: ventral, *L*: worm body length. (B, D) GCaMP dynamics of PDE (*upper*) and the corresponding curvature dynamics from head to tail (*lower*) for freely moving worms (B) and muscularly paralyzed worms restrained within sinusoidal channels (D). (E) Cross-correlation between intracellular calcium dynamics of PDE and curvatures of different body regions from head to tail for freely moving worms (orange, n = 12 animals) and channel-restrained worms (purple, n = 20 animals). (F) Average PDE calcium activity at different values of midbody curvature for freely moving worms (orange) and channel-restrained worms (purple). Curves were obtained via a moving average along the *x*-axis (window size 2), and the filled area represents a 95% confidence interval.

While the finding that PDE calcium activity is correlated with body bending is suggestive of a proprioceptive role for PDE, it does not definitively establish such a function. Other neurons or muscles may influence PDE activity in a manner related to, but not caused by, body bending. To investigate whether PDE activity is caused by body bending, we sought to examine PDE activity in a worm with externally-induced bending and defective muscle contraction. Specifically, we monitored PDE calcium dynamics in *unc-54(e1092)* myosin heavy chain mutants, which are profoundly impaired in muscle contraction. To induce body bending, we developed a sinusoidal-channel microfluidic device and constrained the worms within the channel (Fig. 4C). We then manipulated the worm’s position through the channel by modulating fluid flow to produce different curvature values in the body segments (see *Materials and Methods* for details).

As we displaced the paralyzed worms in the channel, we observed fluctuations in PDE calcium dynamics in response to the induced body posture changes (Fig. 4D). Notably, despite the muscle paralysis of the mutant animals, we observed a strong correlation between PDE fluorescence and the bending curvature of different body segments (Fig. 4E), indicating that body bending alone is sufficient to trigger the neural activity of PDE. We also noticed that the curvature-neuronal activity correlation profiles under paralyzed and freely moving conditions coincided solely in the midbody region (Fig. 4E), suggesting that the midbody might represent the spatial receptive field of the proprioceptive response in PDE neurons. Moreover, our analysis of the relationship between PDE activity and midbody curvature revealed that PDE calcium levels increased as midbody curvature shifted from a dorsal bend to a ventral bend (Fig. 4F), regardless of whether the movement was due to muscle contractions in freely moving worms or external forces from the sinusoidal channels.

Our experiments provide evidence that PDE is capable of proprioceptively responding to midbody curvature. To explore the receptor proteins involved in this process, we screened a comprehensive list of candidate mechanosensitive or proprioceptive receptor genes with expressions in PDE as well as several other mechanosensory receptor genes (Fig. S3) (60, 61). We then asked whether these channels are necessary for CCR. We found that of the mutants tested, only the TRP-2 channel mutant *trp-2(sy691)* displayed a significantly lower CCR than wild-type animals (Fig. S3A). Further investigation, such as conducting PDE-specific rescue of the TRP-2 channel for calcium imaging of PDE and behavioral assays, will be necessary to determine the specific function of the TRP-2 channel in PDE with regard to midbody bending.

### Compensatory curvature response requires the D2-like dopamine receptor DOP-3 in AVK neurons

We next sought to determine what other cellular and molecular components are responsible for CCR downstream of dopamine signaling from PDE neurons.

First, we asked which dopamine receptor(s) were required for CCR. Using the straight microfluidic channel to constrain the midbody of worms, we examined CCR behavior in mutant strains, each lacking a single type of dopamine receptor (DOP-1 through DOP-4; Fig. 5A) and combinations of the DOP-1, DOP-2, and DOP-3 receptors (Fig. 5B). The *dop-3* mutation had a significant negative effect on CCR in all cases tested, and the addition of exogenous dopamine did not restore CCR in *dop-3* mutants. Conversely, dopamine receptor mutants without the *dop-3* mutation displayed normal CCR. We examined the effect of the *dop-3* mutation on CCR induced by optogenetic perturbation and found that *dop-3* mutants again displayed notable defects in CCR induced by midbody muscle inhibition or stimulation (Fig. 5C-E). These results show that the D2-like dopamine receptor DOP-3 is required for CCR.

**Figure 5.**
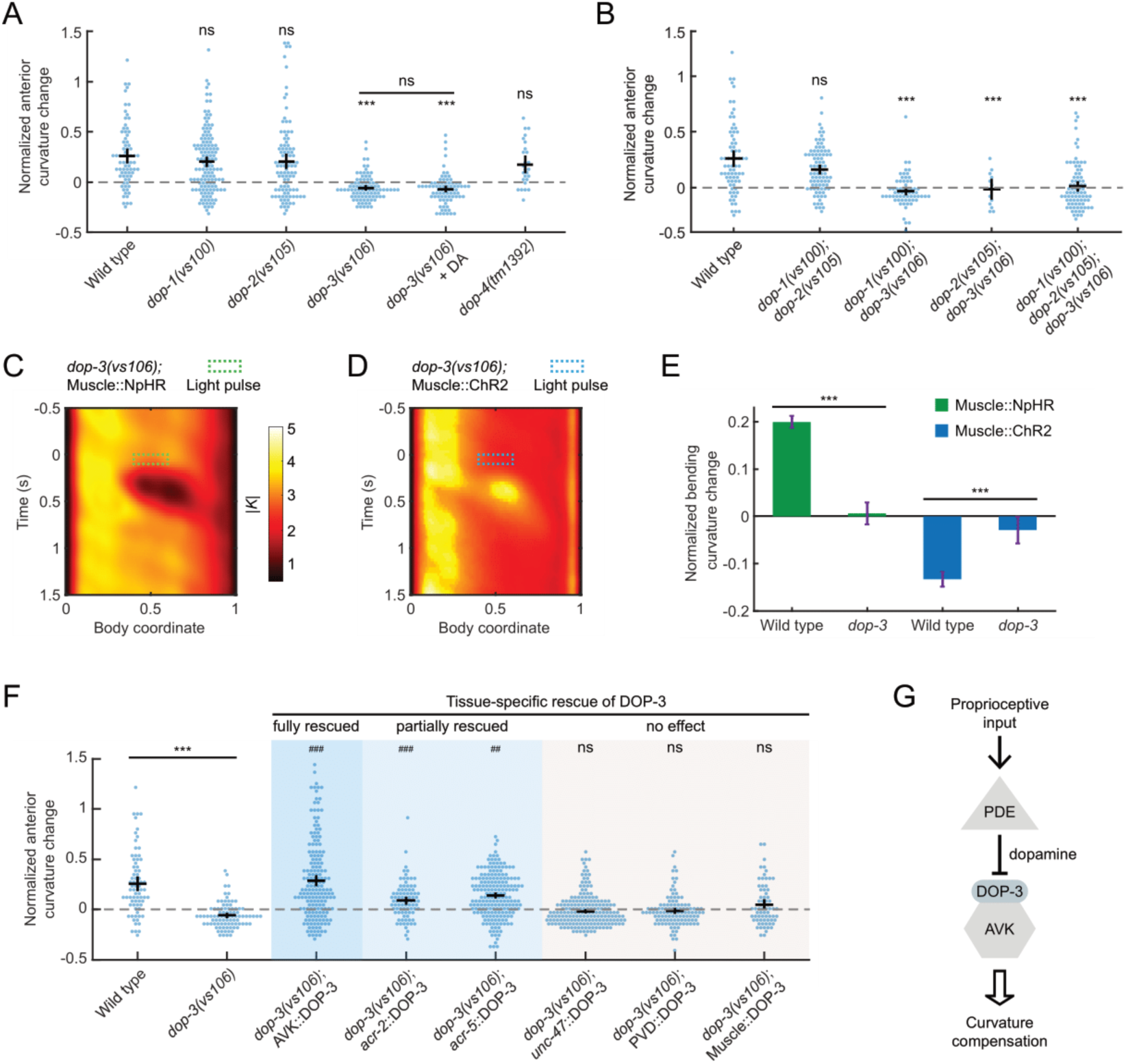
Compensatory curvature response requires the D2-like dopamine receptor DOP-3 in AVK neurons. (A) CCR indices for wild type and dopamine receptor knockout single mutants *dop-1(vs101)*, *dop-2(vs105)*, *dop-3(vs106)*, and *dop-4(tm1392)* under indicated conditions. n ≥ 10 animals per group. Errors bars indicate mean ± SEM. ***p < 0.001 compared with wild type, Dunnett’s multiple comparison tests; ns: not significant when comparing *dop-3* mutants in the absence and presence of exogenous 50 mM dopamine, Student’s t-test. (B) CCR indices for *dop-1 dop-2*, *dop-1 dop-3*, *dop-2 dop-3* double mutants and *dop-1 dop-2 dop-3* triple mutants, compared with wild-type animals. n ≥ 10 animals per indicated condition, mean ± SEM. ***p < 0.001, Dunnett’s multiple comparison tests. (C, D) Mean absolute curvature around 0.1 s illuminations (dashed box) for *dop-3* mutants expressing Muscle::NpHR (C, n = 183 midbody illuminations from 31 worms) or Muscle::ChR2 (D, n = 213 dorsal midbody illuminations from 33 worms). (E) Normalized anterior curvature change of the first post-illumination curvature peak for wild-type and *dop-3* animals expressing Muscle::NpHR (green) or Muscle::ChR2 (blue), mean ± SEM. ***p < 0.001, Student’s t-test. (F) CCR indices for *dop-3* mutants with DOP-3 function rescued by transgenic expression in AVK neurons (Pflp-1(trc)::DOP-3), cholinergic neurons (Pacr-2::DOP-3), B-type motor neurons (Pacr-5::DOP-3), GABAergic neurons (Punc-47::DOP-3), PVD neurons (Pser-2-prom3::DOP-3), and body wall muscle cells (Pmyo-3::DOP-3), compared with wild type and *dop-3(vs106)* mutants. n ≥ 10 animals per condition, mean ± SEM. ***p < 0.001 compared with wild type; ns: not significant, ^###^p < 0.001, ^##^p < 0.01 compared with *dop-3* mutants, Tukey-Kramer multiple comparison tests. See Methods for rescue criteria. (G) A model showing midbody proprioception regulates CCR through DOP-3-dependent dopamine signaling from PDE to AVK neurons.

We next aimed to identify the cell types expressing DOP-3 required for CCR. In wild-type animals, DOP-3 is expressed in several cell types, including GABAergic neurons, cholinergic motor neurons, PVD mechanosensory neurons, AVK interneurons, and body wall muscles (55, 62). Using various promoters (see Table S2), we expressed DOP-3 in these cell types (co-expressed with fluorescent marker on GABAergic neurons via *Punc-47::GFP*) in a *dop-3* mutant background and tested the ability of the transgenes to rescue CCR (Fig. 5F). Restoring DOP-3 expression in GABAergic neurons, PVDs, or body wall muscles failed to rescue the *dop-3* defect (see *Materials and Methods* for rescue criteria). We also examined *dma-1(tm5159)* mutants that lacked the “menorah” structures in PVD dendrites, which are necessary for mechanosensory function (45). This mutant displayed normal CCR (Fig. S3B), suggesting that PVD does not contribute to CCR. However, when we reintroduced DOP-3 expression in AVK neurons (62), CCR was fully restored to a wild-type level (Fig. 5F). A partial rescue of CCR was observed in *dop-3* mutants with DOP-3 expression in cholinergic or B-type motor neurons (Fig. 5F). Our rescue experiments indicate that DOP-3 receptors in AVK and potentially some cholinergic motor neurons mediate the proprioception-triggered dopamine signals from upstream PDE neurons that regulate CCR behavior (Fig. 5G).

We further evaluated CCR in mutants that disrupt the downstream G protein signaling of DOP-3 and DOP-1 (Fig. S3C). Animals with mutations that interfere with the activation of the 𝐺𝛼_*o*_ pathway (linked to DOP-3) exhibited defective CCR, whereas mutants with deficiencies in proteins associated with the 𝐺𝛼_*q*_ pathway (linked to DOP-1) exhibited normal CCR (Fig. S3C; see *Supporting Information*). DOP-3 and DOP-1 elicit opposing effects on locomotion by signaling through these two antagonistic G protein pathways, respectively (55). Our data are consistent with the previously proposed model that DOP-3 affects locomotion by activating the 𝐺𝛼_*o*_Signaling pathway (55) (Fig. S3D) since DOP-3 receptors, but not DOP-1 receptors, were necessary for CCR (Fig. 5A, B),.

These results suggest that CCR requires dopamine signaling through DOP-3 receptors in AVK neurons via 𝐺𝛼_*o*_ pathways.

### The FMRFamide-like neuropeptide FLP-1, released by AVK, regulates SMB motor neurons via receptor NPR-6 to modulate anterior bending amplitude

We have demonstrated that PDE neurons respond to midbody curvature and that the dopamine/DOP-3 signaling pathway from PDE to AVK is required for CCR. The interneuron AVK mediates FLP-1 FMRFamide-like neuropeptide signaling via the release of dense core vesicles (DCVs) to modulate locomotion in response to diverse sensory inputs (42, 62). Deletion of the *flp-1* gene results in loopy undulation with an exaggerated sinusoidal waveform in both agar surface (63) and liquid environments (Movie 4).

We conducted a series of experiments to investigate whether AVK and FLP-1 neuropeptide signaling are necessary for CCR. First, we eliminated AVK neurons by laser ablation (see *Materials and Methods* and *Supporting Information*) and used transgenic animals with AVK expressing tetanus toxin (*Pflp-1::TeTx*) that blocks synaptic vesicle release (62). Both strains showed superficially wild-type locomotion but strongly compromised CCR (Fig. 6A; Movie 5). Next, we tested *flp-1(yn4)* and *flp-1(sy1599)* mutants, which lack the FLP-1 neuropeptide, as well as *unc-31(e169)* mutants, which lack the calcium-activated protein for secretion (CAPS) required for neuropeptide release. We observed significant CCR defects in these mutant animals (Fig. 6B). Transgenic expression of FLP-1 in AVK in *flp-1(yn4)* mutants (62) restored the mutant phenotype (Fig. 6B). These findings demonstrate that FLP-1 neuropeptide signaling from AVK neurons is required for normal CCR.

**Figure 6.**
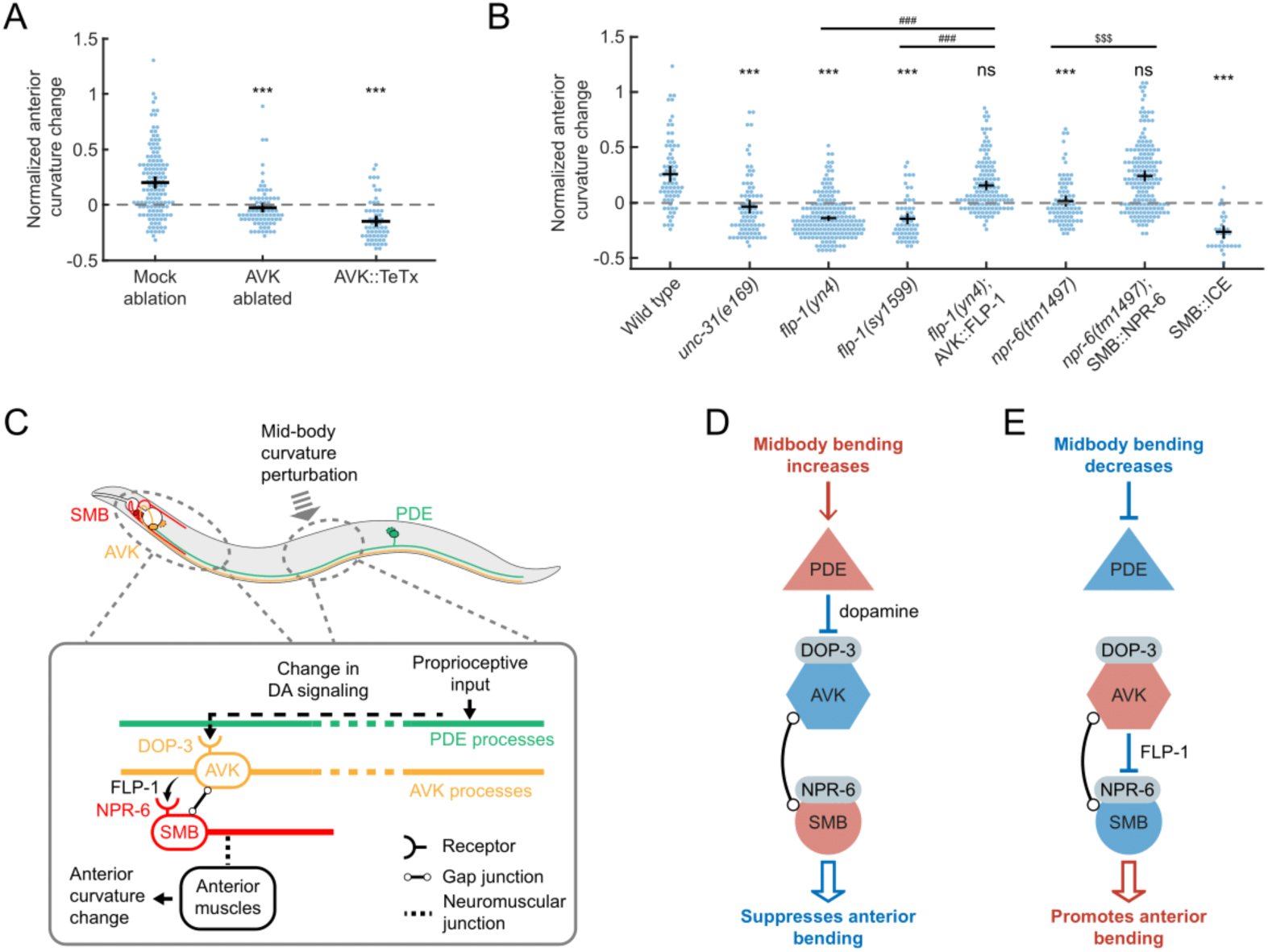
The FMRFamide-like neuropeptide FLP-1, released by AVK, regulates SMB motor neurons via receptor NPR-6 to modulate anterior bending amplitude. (A) CCR indices for animals with laser ablation of AVK and transgenic animals expressing tetanus toxin in AVK (Pflp-1::TeTx), compared with the mock ablation control group. n ≥ 11 animals per condition. Errors bars indicate mean ± SEM. ***p < 0.001, Dunnett’s multiple comparison tests. (B) CCR indices for wild type, *unc-31(e169)*, *flp-1(yn4)*, *flp-1(sy1599)*, *flp-1* mutants with FLP-1 function rescued in AVK, *npr-6(tm1497)*, *npr-6* mutants with NPR-6 function rescued in SMB, and animals lacking SMB (ablation by caspase). n ≥ 12 animals per condition, mean ± SEM. ***p < 0.001, ns: not significant, compared with wild type; ^###^p < 0.001 compared with *flp-1* mutants; ^$$$^p < 0.001 compared with *npr-6* mutants, Tukey-Kramer multiple comparison tests. (C-F) Models for the mechanisms underlying CCR. (C) (*Upper*) Diagram of PDE (green), AVK (orange), and SMB neurons (red) and their somas/processes within a worm body. (*Lower*) Model for CCR. Dopaminergic neurons PDE transduce the proprioceptive input from the midbody curvature and signal to AVK neurons via dopamine signaling through DOP-3 receptors. In the anterior region, the AVK neurons signal via FLP-1 neuropeptides to negatively regulate the head-bending-suppressing motor neurons SMB via NPR-6 receptors. Since PDE negatively regulates AVK via dopamine, AVK negatively regulates SMB via FLP-1 peptides, and SMB negatively regulates head bending, perturbation to the midbody bending leads to a net negative regulatory effect on the anterior bending, as illustrated in two scenarios (D and E). Red and blue indicate excited and inhibited neuronal states, respectively.

The AVK interneurons do not directly innervate muscles to drive body bending. To further probe the circuit underlying CCR, we asked what downstream cells directly affect the anterior bending amplitude while being regulated by FLP-1 signaling from AVK. Previous studies prompted us to speculate that SMB, a class of head motor neurons, might be such a candidate: AVK has both electrical and chemical synapses onto SMB (42), and ablation studies have shown that SMB regulates head and neck muscles and sets the overall amplitude of sinusoidal forward movement (38). Moreover, SMB activity is regulated by AVK-released FLP-1 signaling through the inhibitory receptor NPR-6 (62).

We investigated the role of SMB and its peptide-regulated activity in mediating anterior bending amplitude during CCR. Through the expression of a caspase (62), we ablated SMB neurons, which, in previous studies, led to a significant increase in body bending amplitude (38) (Movie 6). Worms lacking SMBs showed a severely impaired CCR (Fig. 6B). We then examined *npr-6(tm1497)* mutants with and without a transgene that restores the NPR-6 receptor in SMBs (62) and observed restored and defective CCR, respectively (Fig. 6B). These experiments support the idea that the SMB motor neurons modulate anterior bending amplitude under the regulation of AVK-released FLP-1 neuropeptide signaling via the NPR-6 receptor.

Our results, together with the previously reported inhibitory effects of dopamine on AVK and FLP-1 on SMB (42, 62), support the following model for CCR: (1) An increase in midbody bending amplitude promotes dopamine release from PDE, which inhibits FLP-1 release from AVK via the DOP-3 receptor, leading to SMB disinhibition via the NPR-6 receptor and resulting in a decrease in anterior bending amplitude (Fig. 6D); (2) a decreased midbody bending amplitude suppresses dopamine release from PDE, which disinhibits FLP-1 release from AVK and causes SMB inhibition, leading to an increase in anterior bending amplitude (Fig. 6E). This model is also consistent with the observed changes in anterior curvature in response to optogenetic manipulation of PDE neurons (Fig. S4).

### Compensatory curvature response facilitates efficient power expenditure for locomotion

We hypothesized that CCR may help *C. elegans* set its bending amplitude at an optimal level for locomotion in a given environment. *C. elegans* adapts the amplitude, wavelength, and frequency of undulatory movements to environments with a wide range of mechanical loads (5, 6). We recorded animals’ locomotion in viscous fluids and conducted biomechanical analyses for several strains studied for CCR (see *Materials and Methods*). We observed that strains with defective CCR tended to exhibit a larger curvature amplitude compared to strains with normal CCR (Fig. 7A and S6A).

**Figure 7.**
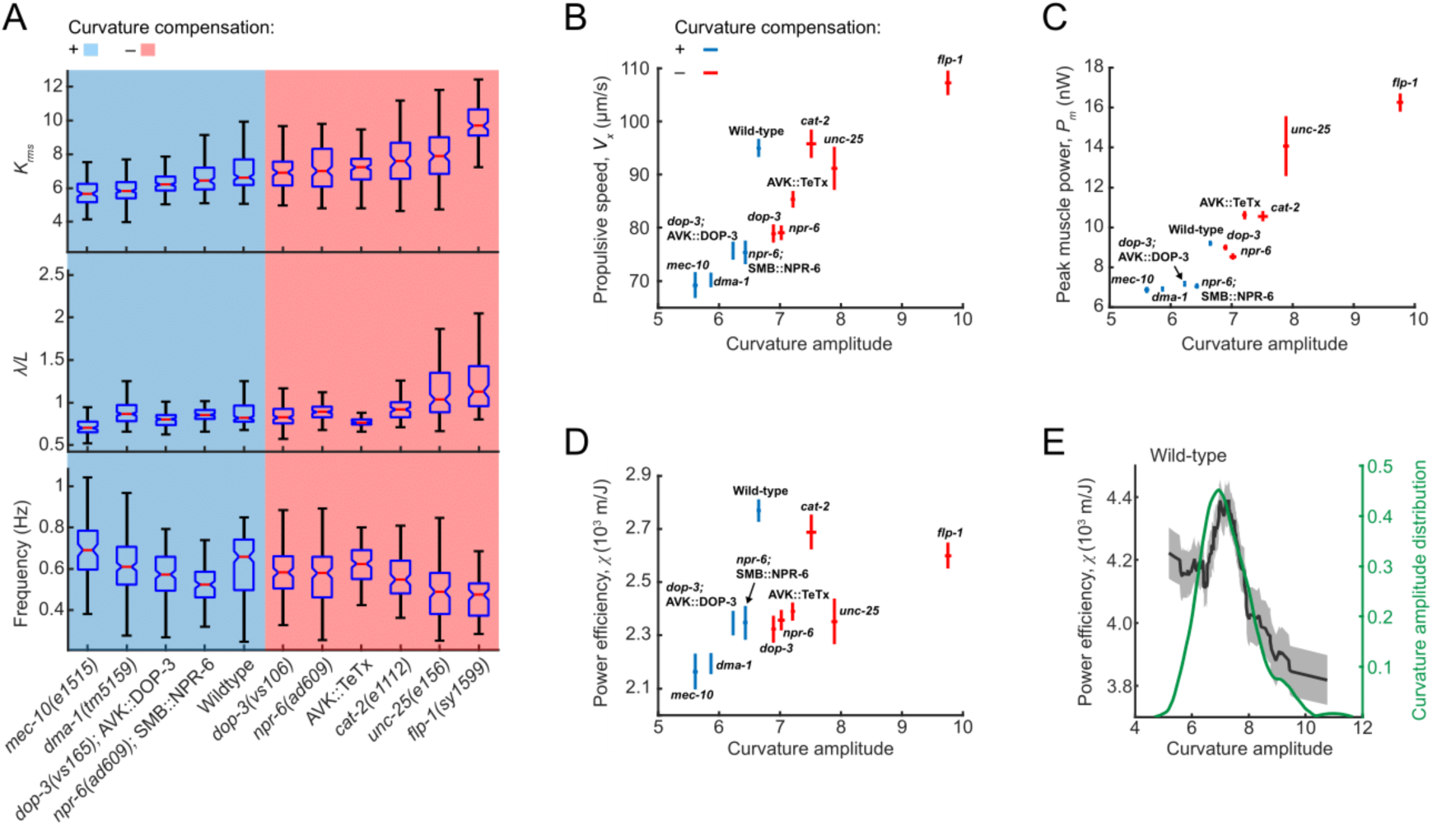
Curvature control promotes efficient power expenditure. (A) Locomotory amplitude, wavelength (scaled by worm body length *L*), and frequency in viscous solutions (120 mPa·s viscosity), measured in selected strains tested for CCR. Blue and red panels denote strain groups with normal and defective CCR, respectively. Each box’s bottom, mid-line, and top represent the data’s 25^th^, 50^th^, and 75^th^ percentiles. The whiskers go from the end of the interquartile range (between the bottom and top of the box) to the 1.5 times that range away from the bottom or top of the box. n = 100-190 forward moving bouts (10 s duration) from 10-20 animals per group. (B) Propulsive speed and curvature amplitude (mean ± SEM), measured from wild-type animals and other strains tested for CCR. Blue and red indicate strains with and without curvature compensation, respectively. (C) Peak muscle power and curvature amplitude (mean ± SEM). (D) Power efficiency and curvature amplitude (mean ± SEM). (E) Power efficiency as a function of curvature amplitude (black) and probability density function of curvature amplitude distribution (green) for wild-type animals (n = 177 forward moving bouts from 30 animals). The 𝜒-𝑲 curve was obtained via a moving average along the *x*-axis (window size 1.5), and the filled area represents a 95% confidence interval within the bin.

Based on direct measurements of locomotion, we defined and quantified the power efficiency of undulatory movements as the ratio of the propulsive speed (Fig. 7B) to the total power required during locomotion (Fig. 7C) (see *Supporting Information* for details). Our results show that compared with the wild-type group, strains with either small or large curvature amplitude displayed lower power efficiency (Fig. 7D).

By examining power efficiency as a function of curvature amplitude in strains with normal CCR, we found that the maximal efficiency was achieved at an intermediate curvature amplitude range consistent with the peak of the curvature amplitude distribution (Fig. 7E and S6B-E). By contrast, in CCR-deficient strains, the peak of the curvature amplitude distribution was not within the optimal amplitude range for power efficiency (Fig. S6F-J).

These results suggest that CCR may provide feedback for optimizing the amplitude of body bending to minimize power usage during forward propulsion. Animals bearing defects in CCR fail to effectively modulate the undulatory amplitude, leading to higher curvature amplitude and reduced locomotor efficiency.

## Discussion

In this study, we have described and characterized a proprioception-mediated compensatory mechanism of locomotor control in *C. elegans*. We demonstrate that, during forward locomotion, the anterior body bending amplitude compensates for the change in midbody bending amplitude by an opposing homeostatic response (Fig. 1H). On the circuit level, we analyzed the sensory and modulatory neuronal components and the signaling molecules required for regulating body posture and undulatory dynamics during the compensatory response to perturbations (Fig. 6C). Our findings provide insights into the neural mechanisms underlying the modulation of undulatory movements in response to proprioceptive cues and highlight the importance of homeostatic control in locomotor behavior.

Our calcium imaging experiments suggest that PDE dopaminergic neurons might function as proprioceptors (Fig. 4). However, further studies are necessary to verify their role in directly detecting midbody movements. Our behavioral analysis indicates that the TRP-2 channel may be responsible for the observed proprioceptive responses in PDE (Fig. S3A) and future *in vivo* calcium imaging of PDE in a *trp-2* mutant background could help confirm this role. Additionally, investigating the expression of TRP-2 gene may help localize the proprioceptive components in PDE.

To clarify the neurotransmitter systems involved in CCR, we tested dopamine-deficient mutants and their response to exogenous dopamine. Previous studies have used exogenous dopamine to restore other *C. elegans* dopamine-related behaviors, such as basal slowing response (50), area-restricted search (48), and gravitaxis (64), but without distinguishing between the roles of dopamine during behavioral assays or development. Unlike these behaviors, CCR in *cat-2* mutants was restored only by exogenous dopamine administered shortly before the assay (about 3 min), indicating an active role for dopamine in CCR competence. In addition, the failure of added dopamine to rescue CCR in animals lacking or with defective synaptic transmission in PDEs (Fig. 3H) suggests that CCR requires phasic dopamine transmission, specifically from PDE.

Our experiments show that DOP-3 is required for CCR, and the restoration of CCR by genetic DOP-3 expression in AVK neurons confirms their role in mediating the dopamine effect on CCR. Partial restoration was observed from genetic DOP-3 expression in cholinergic or B-type motor neurons. The reason for these partial rescues is currently unclear, but it may result from a potential AVK expression under these promoters, as indicated by gene expression dataset (60). Alternatively, these motor neurons might constitute a parallel circuit acting redundantly to mediate CCR.

Downstream of the PDE dopamine signaling, the CCR circuit comprises the FLP-1 FMRFamide-like neuropeptide signaling from AVK to the head motor neurons SMB that regulate the sinusoidal amplitude (38). This circuit overlaps with the one for a food-induced behavior (62), where PDE functions as a mechanoreceptor for food presence, and SMB integrates signals from AVK and DVA that antagonistically affect locomotion. AVK is involved in multiple sensory modules, including food detection (62), oxygen sensation (42), chemotaxis (65), and proprioception (this work). Understanding how these sensory cues are integrated into the PDE-AVK-SMB neuronal module would shed light on the neural network controlling behaviors induced by distinct sensations and the complexity within individual neurons and circuits (66).

Proprioceptive feedback is essential for sustaining normal locomotion in *C. elegans* (19), directly transmitting rhythmic bending activity from anterior to posterior segments during forward locomotion (52). However, the proprioceptive information necessary for CCR is not required for normal locomotion, as most animals with defective CCR can still perform regular locomotion, although the anterior curvature amplitude may differ across strains (Fig. S6A). To investigate the effect of CCR on locomotion in viscous fluids, we conducted biomechanical measurements to estimate the energy expenditure during forward locomotion, and our analysis indicates that CCR dynamically regulates bending amplitude within an optimal range for efficient locomotion.

The optimal locomotor efficiency observed under our off-food conditions could be important for *C. elegans* exploratory foraging behavior. While on food bacteria, it exhibits dwelling behavior characterized by slow forward movements and frequent reversals (50). In contrast, when deprived of food, it switches to roaming behavior, which includes long forward runs and infrequent reorientations (38, 48, 59). CCR could thus minimize energy expenditure during food search.

Our study uncovers the ability of *C. elegans* to modulate and maintain the ongoing locomotion based on local bending amplitude. Such a form of locomotor homeostasis highlights the behavioral plasticity of its compact nervous system. A recent study in adult zebrafish reported a similar motor control mechanism, where inhibitory feedback signals modulate locomotor movements based on local mechanical tension (20), indicating a potential conservation of this behavior across species. The identified neuronal and genetic pathways in *C. elegans* may serve as a guide for future studies of motor control in other organismal systems.

## Materials and Methods

### Strains, Cultivation, and Ablation Methods

*C. elegans* were grown at 20°C on nematode growth media (NGM) plates with OP50 *Escherichia coli* using standard methods (67). For optogenetic experiments, animals were raised in the dark on OP50-ATR plates, made by mixing 2 µm of 100 mM all-*trans* retinal (ATR) in ethanol into a 250 µL suspension of OP50 in LB medium and seeded onto 6 cm NGM plates. All experiments were performed with 1-day-old adult hermaphrodites synchronized by hypochlorite bleaching. The strains used and the procedure for constructing plasmid and generating transgenic strains are described in *Supporting Information*. Laser ablation was performed using a previously described thermal laser ablation system (58); for details see *Supporting Information* .

### Behavioral Assays

Optogenetic manipulation experiments employed a laser targeting system adapted from previous methods (19, 53). Microfluidic experiments were conducted using straight and sinusoidal-channel microfluidic devices for CCR assay and calcium imaging of PDE neurons, respectively, on custom high-resolution imaging stages. Kinematic and ethological analyses of animal locomotion followed previous studies (6, 33). Details are available in *Supporting Information* for apparatus configurations, media preparation, data quantification, biomechanical analyses, and statistical analysis.

### Calcium Imaging

Calcium imaging adapted previously described protocols for measuring the activity of the PDE neurons (41). Imaging optics and procedures for preparing and manipulating animals is described in *Supporting Information*. Data analysis was performed using custom software written in MATLAB (MathWorks) and Python.

## Acknowledgments

We thank Niels Ringstad, Kang Shen, Steven Flavell, Andres Villu Maricq, Mei Zhen, Cori Bargmann, Alexander Gottschalk, and Michael Koelle for providing strains and plasmids. Some strains were provided by the *C. elegans* Genetic Center, funded by the NIH Office of Research Infrastructure Programs (P40 OD010440). We thank Gal Haspel, David Raizen, Julia Raizen, Niels Ringstad, Yen-Chih Chen, and Michael Nusbaum for helpful discussions and suggestions. This work was supported by the National Institutes of Health (R01NS084835).

## Figures and Tables

## Supporting Information for

A proprioceptive feedback circuit drives *C. elegans* locomotor adaptation through dopamine signaling

## SI Experimental Results

### CCR analysis of mutants with disruptions in G protein signaling of DOP-3 and DOP-1

We examined the role of downstream G protein signaling in CCR using mutants affecting the 𝑮𝜶_𝒐_ and 𝑮𝜶_𝒒_ pathways (Fig. S3C, D). Mutants with a deficiency in the *C. elegans* ortholog of 𝑮𝜶_𝒐_, GOA-1 (coupled to DOP-3), and in the putative downstream effector of GOA-1, the RGS protein DGK-1, exhibited defective CCR.

In contrast, mutants lacking EGL-10, which inhibits GOA-1 𝑮𝜶_𝒐_, and GPB-2, a subunit of EGL-10 RGS, exhibited wild-type CCR. Mutants deficient in 𝐺𝛼_*q*_-associated proteins (coupled to DOP-1), including EGL-30, EGL-8, and EAT-16, also showed normal CCR.

### SI Materials and Methods

#### Strains and Plasmids

Bristol N2 was used as the wild-type strain. The complete strain list and plasmid list used in this study are available in Tables S1 and S2, respectively.

To rescue *dop-3* function in specific tissues, transgenic strains were generated via microinjection of plasmid DNA and a fluorescent co-injection marker. Plasmid constructs for tissue-specific expression of DOP-3 include pYX36(*ser-2-prom3::dop-3*), pYX37(*myo-3::dop-3*), pYX38(*acr-2::dop-3*), pYX39(*acr-5::dop-3*). The *Pser-2-prom3* (PVD), *Pmyo-3* (body-wall muscles), *Pacr-2* (cholinergic neurons), and *Pacr-5* (B-type motor neurons) promoters were constructed from donor plasmids *Pser-2prom3::GFP* (gift from Kang Shen), *Pmyo-3::RCaMP1h* (gift from Alexander Gottschalk), *Pacr-2::GFP* (gift from Michael Koelle), and *Pacr-5::Arch::GFP* (gift from Shin Takagi), respectively. *dop-3* gene sequence was amplified from pDC50(*unc-47::dop-3*) using primers SAZ86 and SAZ87. Constructs containing promoter sequences were amplified from the corresponding donor plasmids using primers SAZ88 and SAZ89. Reconstruction procedures were conducted using the Gibson Assembly method (Gibson Assembly Master Mix, New England Biolabs). The resulting plasmids were verified by sequencing (ABI 3730XL sequencer, Penn Genomic Analysis Core).

#### Laser Ablation

Using an infrared laser ablation system (1), we performed cell ablation experiments on transgenic animals (*Pdat-1::GFP* for PDE ablation; *Pflp-1::GFP* for AVK ablation) at the third larval stage. Worms were immobilized on 10% agar pads with 50 nm polystyrene beads (2). Under GFP fluorescence optics, the target neurons (PDE or AVK) were illuminated with 1∼2 laser pulses (1.5 ms duration, 400 mW power) through a 63X oil-immersion objective. Animals were transferred to a fresh OP50 plate after ablation to recover overnight and then re-mounted to the system to confirm the elimination of target neurons. Confirmed animals were transferred back to seeded plates to grow into young adults. For the PDE-vicinity ablated group, we lesioned each worm proximal to PDEs (about 5 µm away from the center of the GFP-labeled cell soma) without damaging the PDE somas. Mock controls were also mounted but not irradiated with the laser.

#### Optogenetic Experiments

For experiments with optogenetic manipulation, worms were prepared in a viscous solution using 17% (by mass) dextran (Sigma-Aldrich D5376, average molecular weight 1,500,000-2,800,000) in NGM buffer (120 mPa·s viscosity) confined within chambers formed by a microscope slide and a coverslip separated by 125-µm-thick polyester shims (9513K42, McMaster-Carr).

We used a Leica DMI4000B microscope with a motorized stage to conduct optogenetic experiments, recording image sequences at 50 Hz with an sCMOS camera (optiMOS, Photometrics) at 10X magnification (Leica Plan Fluotar; N.A. 0.30) under dark field illumination from red LEDs. Using a custom-built optogenetic targeting system (3), we optogenetically manipulated worm muscle or neural activity during locomotion with spatial selectivity. Inhibition or stimulation of muscles/neurons was achieved with a green (532 nm wavelength) solid-state laser (GL532T3-300, Shanghai Laser & Optics Century) at 16 mW/mm^2^ or a blue (473 nm wavelength) solid-state laser (BL473T3-150, SLOC) at 10.3 mW/mm^2^.

For optogenetic inhibition and stimulation of PDE neurons, we used transgenic animals with all dopaminergic neurons expressing inhibitory opsin *GtACR* or excitatory opsin *CoChR*, respectively (via *Pdat-1*). During experiments, each animal was illuminated at 0.4-0.8 body region by a brief laser pulse (with stated durations) repeated 10 times, with a 6 s interval between pulses.

For optogenetic muscle inhibition and stimulation, we used wild-type and mutant animals with body wall muscles expressing inhibitory opsin *NpHR* or excitatory opsin *ChR2*, respectively (via *Pmyo-3*). During experiments, each animal was illuminated at indicated regions (both sides for inhibition, dorsal or ventral side for stimulation) by a brief laser pulse (0.1 s duration, unless otherwise stated) repeated 10 times, with a 6 s interval between pulses.

For experiments with modulated optogenetic manipulation of muscles, we used pulse durations of 0.1-0.5 s and modulated the irradiance of laser pulses delivered to animals (3-16 mW/mm^2^ for the 532 nm laser, 2.5-10 mW/mm^2^ for the 473 nm laser). To vary laser power, we placed two optical polarizers (Vivitar, Thorlabs) in the optical pathway and adjusted the output light power by rotating the first polarizer.

#### Microfluidic Experiments

We fabricated two types of custom microfluidic polydimethylsiloxane (PDMS) devices using standard soft lithography techniques (4).

The straight-channel microfluidic device consists of 2000-µm-wide open areas connected by two parallel straight channels (width x length x height: 60 µm x 200 µm x 80 µm). The microfluidic chamber was loaded with NGM buffer containing 0.1% (by mass) bovine serum albumin (BSA) to prevent worms’ adherence to chamber surfaces or tubing. The sinusoidal-channel microfluidic device consists of a 60-µm-wide sinusoidal channel with a sine wave measuring 200 µm in height and 400 µm in wavelength. The chamber of this device was loaded with NGM buffer mixed with 0.1% BSA and 17% (by mass) dextran. In both types of microfluidics, we use polyethylene tubing (Saint-Gobain) and a 3-way Luer valve (Cole-Parmer) to connect the chamber of the microfluidic device parallel to a 1 mL syringe and an NGM buffer reservoir. We finely adjusted the flow by rotating a screw that compressed the tubing slightly between the chamber and the syringe.

We used straight-channel microfluidics to perform the CCR assay. First, we transferred young adults to a food-free NGM buffer for about 5 min to remove any bacteria carried by the animals. Next, we pipetted the animals from the buffer into the inlet of the microfluidic chamber. To move worms to the field of view (approximately 4 mm x 3 mm), we applied pressure or vacuum using the syringe on the inlet. We recorded behavioral videos of each animal in the field of view for 3 minutes, including periods of both constrained and free locomotion.

Video sequences were recorded at 30 frames per second with a 5-megapixel CMOS camera (DMK33GP031, The Imaging Source) and a C-mount lens (Nippon Kogaku NIKKOR-H; effective focal length, 28 mm) using IC Capture software (The Imaging Source). Dark-field illumination was provided by red LED rings (outer size, 80 mm; Qasim) surrounding the device.

In the CCR assays conducted under different food conditions (Fig. S2C), we initially inoculated a 50-ml lysogeny broth (LB) medium culture with OP50 and allowed it to grow for 24 h at 20°C. We then centrifuged an aliquot of this culture and resuspended it with NGM buffer at concentrations 0.125x and 1x that of the original liquid culture (5). For CCR assays in the absence of food, we used food-free NGMB solutions.

We used sinusoidal-channel microfluidic devices to record PDE calcium activity in animals subjected to externally-applied bending. Methods for worm loading and position manipulation were similar to those used in the straight-channel microfluidic device.

#### Behavioral Data Quantification

The behavioral data from optogenetic and microfluidic experiments were processed using MATLAB custom software (MathWorks) as described in previous reports (3, 6). The normalized curvature 𝐾 is the product of 𝜅 and the worm body length 𝐿, derived from the worm centerline length. We excluded the anterior and posterior 5% body regions from the analysis to avoid high-frequency movements. The moving direction of a worm was determined by the gradients in the curvature over time and body coordinate, and image sequences during which the worm moved forward for at least 4 seconds were selected for analysis. The curvature dynamics of the anterior, middle, and posterior regions were defined as the average of the normalized curvature over 0.1-0.3, 0.4-0.6, and 0.7-0.9 body coordinates, respectively.

To quantify the effect of optogenetic perturbations on worm undulatory amplitude, we calculated the curvature amplitude of the anterior and middle regions around each trial of laser illumination. We used the MATLAB function *findpeaks* to identify local extrema along the time-varying curvature profiles. We quantified the change in curvature amplitude induced by optogenetic perturbations using the corresponding normalized curvature change defined by 𝛥𝐾_+𝑛_/|𝐾_−𝑖_| = (|𝐾_+𝑛_| − |𝐾_−𝑖_|)/|𝐾_−𝑖_|, where 𝐾_)*_ denotes the value of the 𝑛^th^ post-illumination curvature peak, and 𝐾_+,_ (𝑖 = 1 or 2) denotes the value of the 𝑖^-.^ last pre-illumination curvature peak with the same sign as 𝐾_)*_.

To quantify the effect of straight-channel constraint on worm undulatory amplitude, the whole-body curvature amplitude during constrained locomotion was computed and compared with free locomotion.

We analyzed worm locomotor dynamics during free locomotion to generate an averaged curvature dynamics profile as a function of body coordinates. We divided the worm body coordinate into 10 sections from head to tail (from 0.05 to 0.95, omitting the anterior and posterior 5% regions) and calculated the average of the normalized curvature over the body coordinate of the section for all periods of free locomotion. Local extrema along each time sequence of curvature were identified using the *findpeaks* function in MATLAB, and the mean of the absolute value of these local extrema was used to denote the curvature amplitude at the body coordinate defined by the midpoint of the section (e.g., 0.1 for section 0.05-0.15). After computing curvature amplitudes for the 10 sections, we obtained the whole-body averaged curvature amplitude profile, 𝐴_/011_(𝑠), through a linear 1-D interpolation with 100 sample points of values computed across the worm body.

We divided video sequences of constrained movement into individual short sequences using a 3 s time window to analyze constrained locomotion. Due to fluctuations in the worm position controlled by the syringe, the constrained body region did not remain fixed. To record the relative position of the constraint to the worm body (gray lines in Fig. S2B), we manually marked the channel position in each image sequence.

The normalized curvature change in response to mid-body constraint was calculated by considering only periods during which the anterior and posterior limits of the narrow channel were consistently within 0.35-0.65 body coordinates. The resulting curvature dynamics were denoted as *K_const_*, and the maximum value of ⃒*K_const_*(*s,t*)⃒ in the time dimension for all qualified short periods was defined as the curvature amplitude profile of individual periods, denoted as *A_const_*(*s*) = max/t⃒*K_const_*(*s,t*)⃒. The normalized curvature change of each period was represented *A_const_*(*s*)*A_free_*(*s*) − 1. The normalized anterior, mid-body, and posterior curvature changes of individual periods were 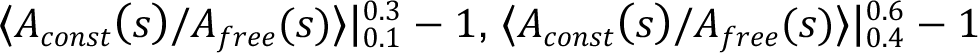, and 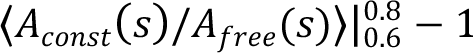, respectively, where 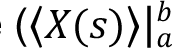 denotes the average of 𝑋 over the interval [a, b].

#### Calcium Imaging

The method for recording PDE calcium activity differed for freely behaving and paralyzed animals. For freely behaving animals, transgenic worms expressing GCaMP in dopaminergic neurons were placed on a 5% agar pad topped with 2 µL NGM buffer and covered with a #1.5 cover glass. Worms moved slowly under this condition while maintaining normal body shape and locomotion. For paralyzed animals, *unc-54(e1092)* myosin heavy chain mutants expressing GCaMP in PDE neurons were loaded into a sinusoidal-channel microfluidic device filled with 17% dextran solution. The worm position within the channel was controlled manually using a 1 mL syringe to induce varying body curvature.

Calcium imaging of PDE in both freely behaving and paralyzed worms was performed on a Leica DMI3000B microscope with a motorized stage (CTR3000, Leica). GCaMP6m protein in PDE neurons was excited by broadband excitation light from a Leica EL6000 illuminator. Through a red filter, we visualized the worm body using a built-in halogen lamp (LH107/2, Leica). To enable simultaneous recording of calcium activity and worm movement, we collected green fluorescence emission and red, dark field illumination using a Leica Plan Apo 10X objective (N.A. 0.40), separated by a dual-view beam splitter (DV2, Photometrics) with a GFP/RFP filter set, projected onto an EMCCD sensor (Cascade 1K, Photometrics). We acquired image sequences at 9 frames per second (fps) with 100 ms exposure time, using MicroManager software, and acquired approximately 2 min of data per animal.

We used custom analysis routines to process the acquired image sequences from both worm preparations. Each image in the sequence was split in half to separate the red and green channels and then background subtracted and thresholded to generate binary images in the red channel. The binary image sequences were used to quantify worm curvature dynamics, then to computationally deform the worm outline into a straightened rectangle. This mask was then used to crop out whole-body fluorescence signals from the green channel, and regions of interest (ROIs) were selected on somas of PDE neurons. We measured GCaMP signals as integrated fluorescence over the ROI, with background subtraction from a secondary ROI drawn around PDEs but lacking labeled neurons. Normalized signals were obtained by subtracting and dividing GCaMP values by the baseline 𝐹_D_ value calculated per recording as the mean of the lowest 50% GCaMP values. We used ImageJ software (Fiji, ImageJ2) for image splitting, binarization, and GCaMP signal extraction, and custom-written Python scripts for curvature calculation and binary image deformation.

#### Quantification and Statistical Analysis

Each transgenic and mutant strain was tested in at least two experiments performed on two separate days within a week and compared to control experiments run in parallel on the same days. All quantification details are provided in *Supporting Information*. We reported the specifications of all statistical analyses in the figure legends. We considered differences significant if p < 0.05. For DOP-3 rescue experiments, we considered DOP-3 expression in a specific tissue as a full rescue for the *dop-3* mutant phenotype if the data were not significantly different (p > 0.05) from wild-type animals assayed in parallel. We considered it a partial rescue if the data were significantly greater (p < 0.05) than that of *dop-3* mutants and significantly smaller (p < 0.05) than that of wild-type animals. We considered it having no effect if the data were not significantly different from *dop-3* mutants.

#### Kinematic and Ethological Analyses of Locomotion

We calculated the undulatory wavelength 𝜆 and frequency 𝑓 by analyzing curvature, using previously described methods (7, 8). We quantified the forward moving speed 𝑉_E_ using the resistance force theory of slender swimmer (9). Specifically, we used the expression 𝑉_E_ = 𝛼_3_𝑉_F_, where 𝑉_F_ = 𝑓 · 𝜆 is the backward velocity of the undulatory waves relative to the worm body, and 𝛼_G_ = (𝐶_H_/𝐶_I_ − 1)/(2𝜃^+K^ + 𝐶_H_/𝐶_I_) is a nondimensional coefficient depending only on the ratio between the normal and longitudinal drag coefficients 𝐶_H_/𝐶_I_ and the peak angle of attack 𝜃_J_ of a body segment with respect to the direction of motion (6). The ratio of drag coefficients 𝐶_H_/𝐶_I_ was set to 1.6 according to previous experimental and theoretical results (8, 9). The muscle power 𝑃_L_ required for undulations was given by (6):

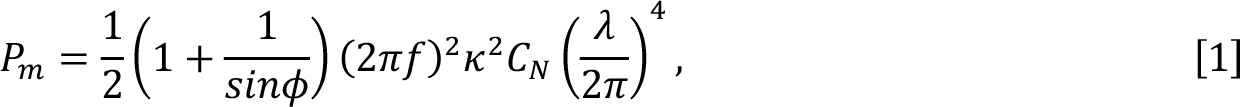

where 𝜙 =𝑎𝑟𝑐𝑡𝑎𝑛 (2𝜋𝑓𝐶_H_𝜆^M^/𝑏(2𝜋)^M^) is the phase difference between curvature and the bending torque produced by muscles, and 𝐶_H_ is estimated to be 4587 mPa·s, based on 3.3 times the fluid viscosity (7). Among all strains tested, we observed a positive correlation between both 𝑉_E_ and 𝑃_L_ with curvature amplitude (Fig. 7B, C).

We defined the power efficiency 𝜒 as the ratio of the propulsive speed 𝑉_E_ to the sum of the muscle power 𝑃_L_ and the basal metabolic power 𝑃_N_, where 𝑃_N_ was set to 60 nW based on reported values from oxygen consumption rate measurements (10).

**Fig. S1.**
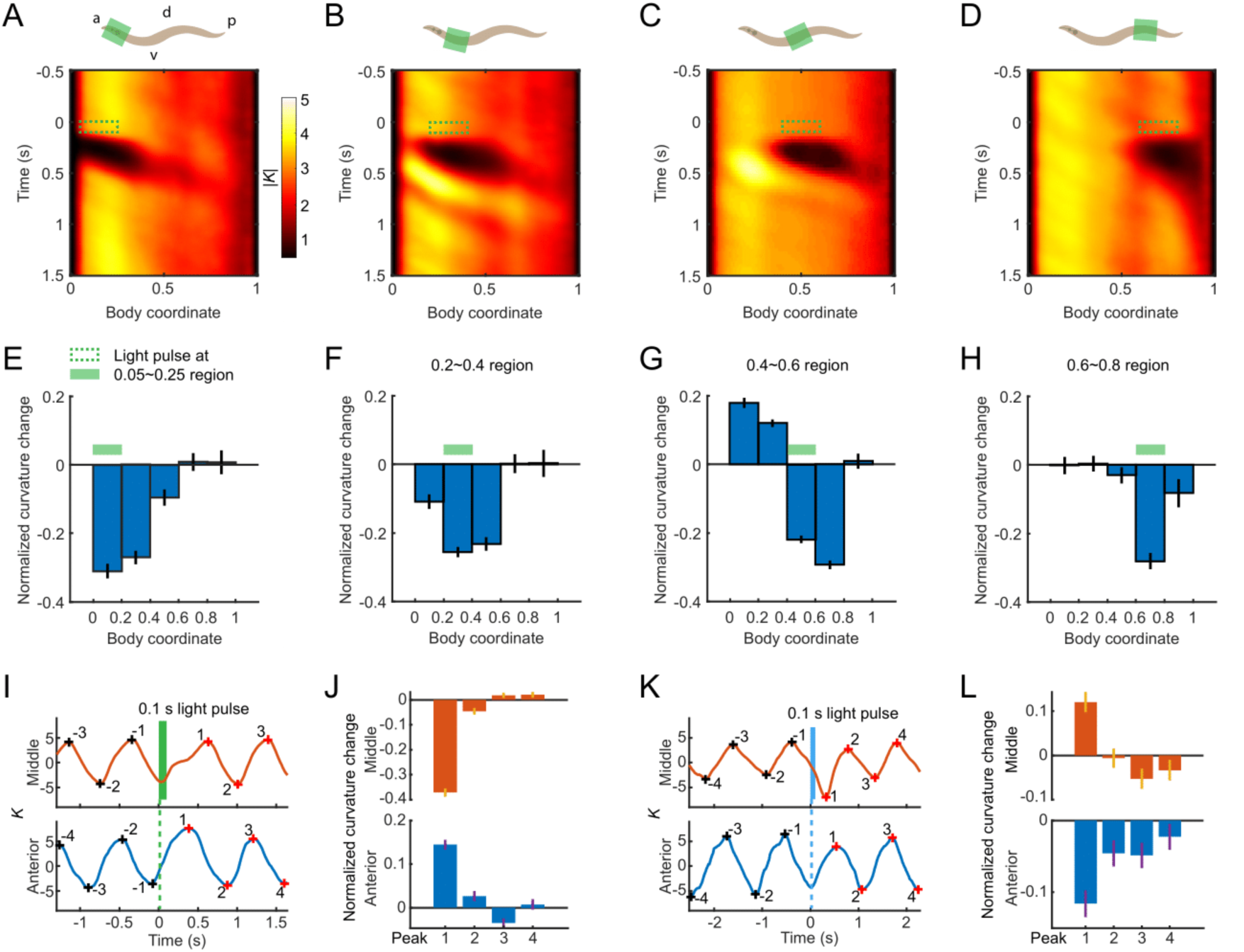
Curvature modulation in response to optogenetic muscle inhibition at various body regions. (A-D) (*Upper*) Optogenetic muscle inhibition (green bar) applied at head (A), neck (B), middle (C), and posterior (D) regions of transgenic worms expressing Muscle::NpHR. a: anterior, p: posterior, d: dorsal, v: ventral. (*Lower*) Mean absolute curvature. Green box indicates laser illumination. 649 illuminations from 135 worms for (A); 466 illuminations from 75 worms for (B); 1160 illuminations from 206 worms for (C); 467 illuminations from 76 worms for (D). (E-H) Undulatory amplitude change upon transient optogenetic muscle inhibitions applied at indicated body regions, measured as mean ± SEM of the normalized curvature change of the first post-illumination curvature peak of various body regions from the head to the tail. Green bar indicates a 0.1 s laser illumination applied to the corresponding body region. (I-K) Curvature dynamics of a worm’s middle (*upper*) and anterior (*lower*) regions around a 0.1 s muscle inhibition (I, green) or stimulation (K, blue) in the mid-body. Black and red crosses mark the last 4 pre-illumination and the first 4 post-illumination curvature peaks, respectively. (J-L) Undulatory amplitude change upon transient mid-body muscle inhibitions (J, n = 1160 mid-body illuminations from 206 worms) or stimulations (L, n = 693 dorsal mid-body illuminations from 122 worms), measured as mean ± SEM of the normalized curvature change of the first 4 post-illumination curvature peaks of the middle (*upper*) and anterior (*lower*) body regions.

**Fig. S2.**
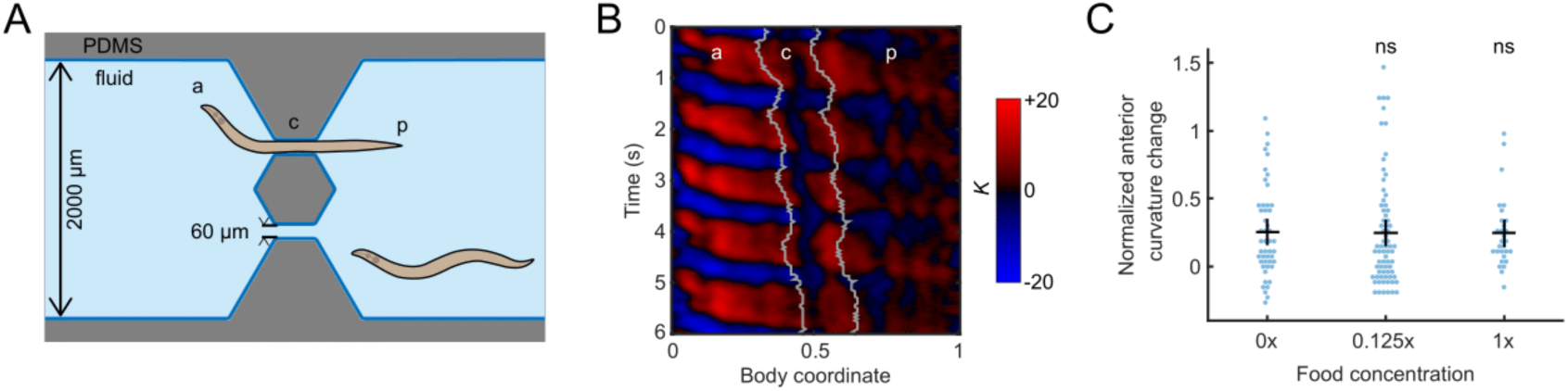
Microfluidics for physically constraining mid-body bending movements during locomotion. (A) Straight-channel microfluidic device for constraining body curvature. a: anterior region, p: posterior region, c: constrained middle region of a worm. (B) Curvature of a bout of constrained movements. Gray lines indicate the anterior and posterior limits of the constriction. (C) CCR indices for wild-type animals under different food conditions (see *Supporting Information*). n ≥ 10 animals per group, mean ± SEM. ns: not significant compared to the group under food-free conditions, Dunnett’s multiple comparison tests.

**Fig. S3.**
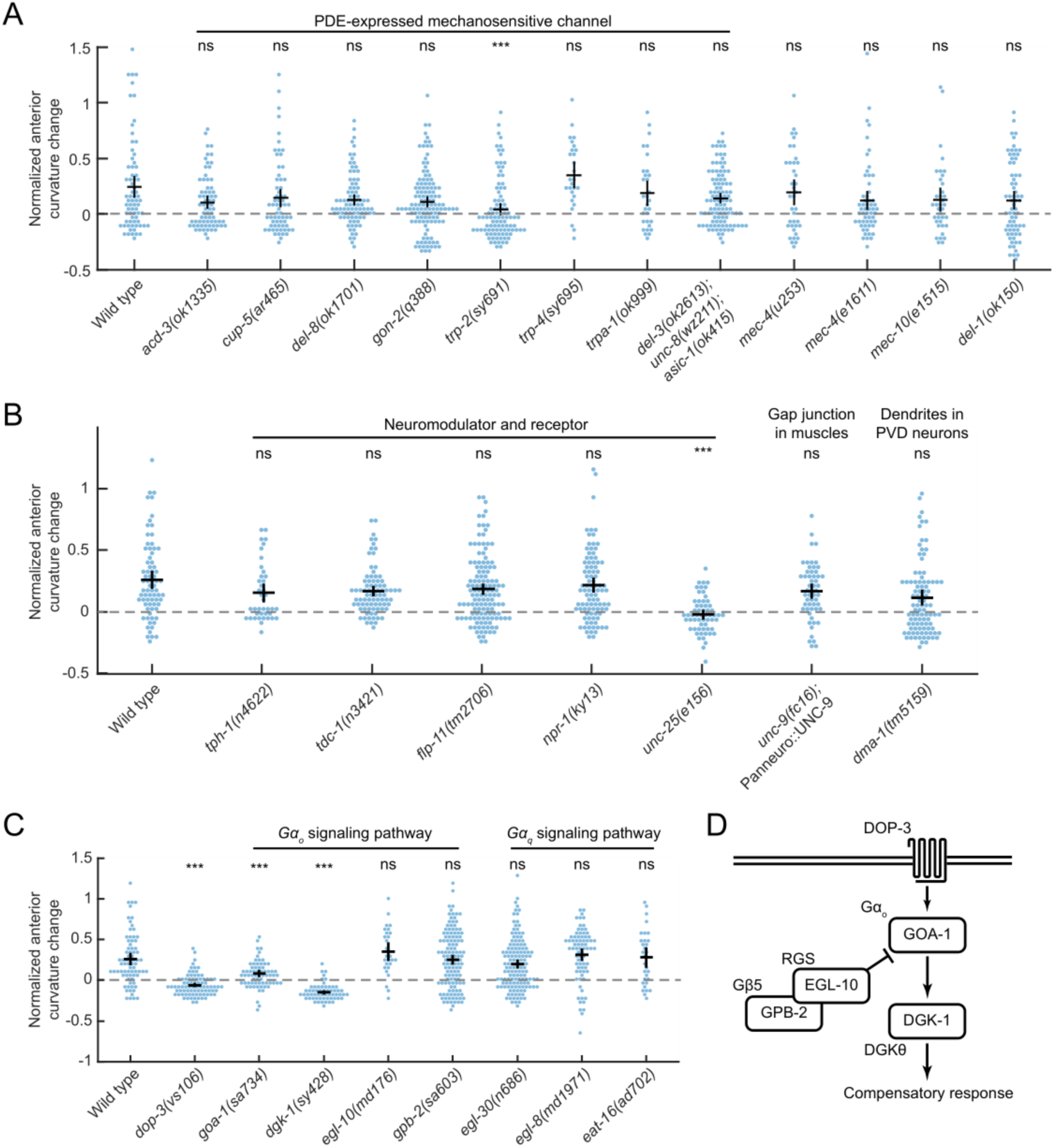
Genetic candidate screening of additional strains for CCR. (A) CCR indices for PDE-expressed mechanoreceptor mutants *acd-3(ok1335)*, *cup-5(ar465)*, *del-8(ok1701)*, *gon-2(q388)*, *trp-2(sy691)*, *trp-4(sy695), trpa-1(ok999),* and triple mutant *del-3(ok2613)unc-8(wz211)asic-1(ok415)*, and non-PDE-expressed mechanoreceptor mutants *mec-4(u253)*, *mec-4(e1611)*, *mec-10(e1515)*, *del-1(ok150)*. n ≥ 10 animals per group, mean ± SEM. ***p < 0.001, ns: not significant compared with wild-type animals, Dunnett’s multiple comparison tests. (B) CCR indices for *tph-1(n4622)*, *tdc-1(n3421)*, *npr-1(ky13)*, *unc-25(e156)*, *dma-1(tm5159)* mutants, and *unc-9(fc16)* mutants in which the UNC-9 innexin protein was rescued pan-neuronally. The *unc-9* rescued strain lacks gap junction only in muscle cells. DMA-1 is required to grow the 2°, 3°, and 4° dendrites in the PVD neurons (11). n ≥ 10 animals per group, mean ± SEM. ***p < 0.001, ns: not significant compared with wild-type animals, Dunnett’s multiple tests. (C) CCR indices for *dop-3(vs106)* mutants, 𝑮𝜶_𝒐_ signaling mutants *goa-1(sa734)*, *dgk-1(sy428)*, *egl-10(md176)*, *gpb-2(sa603)*, 𝑮𝜶_𝒒_ signaling mutants *egl-30(n686)*, *egl-8(md1971)*, *eat-16(ad702)*, compared with wild-type animals. n ≥ 10 animals per group, mean ± SEM. ***p < 0.001, ns: not significant, Dunnett’s multiple comparison tests. (D) Schematic representation of the 𝑮𝜶_𝒐_ protein signaling pathways that regulate the curvature compensatory response in *C. elegans*. The diagram is based on Chase et al. (2004) (12).

**Fig. S4.**
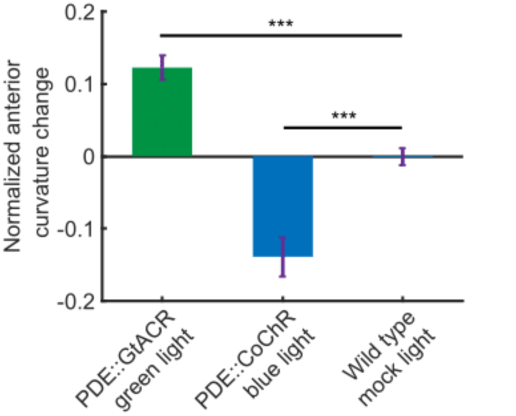
*C. elegans* modulates anterior bending amplitude in response to optogenetic manipulation of PDE neurons. Normalized anterior curvature change for the first post-illumination curvature peak for transgenic animals expressing P*dat-1*::GtACR (green, n = 50 worms) illuminated by 0.3 s green light and transgenic animals expressing P*dat-1*::CoChR (blue, n = 12 worms) illuminated by 0.5 s blue light, compared to the wild-type control group (mock light: no light during illumination events, n = 116 worms). All animals were illuminated at 0.4-0.8 body coordinate. Error bars indicate mean ± SEM. ***p < 0.001, Dunnett’s multiple comparison tests.

**Fig. S5.**
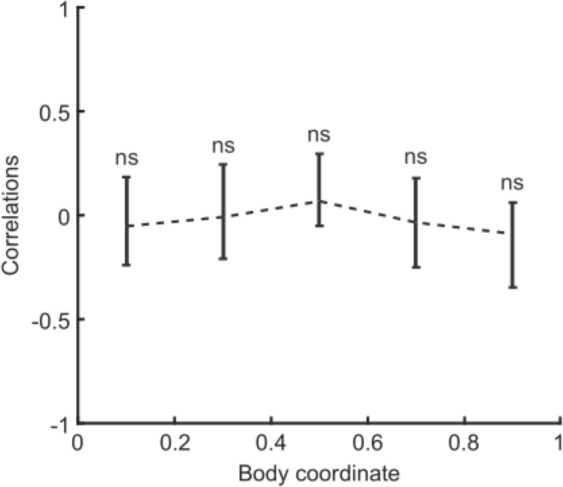
Correlation coefficient of PDE::GFP signal with body curvature of various body regions for freely moving worms. N = 15 animals, mean ± SEM. Ns: not significant compared with zero.

**Fig. S6.**
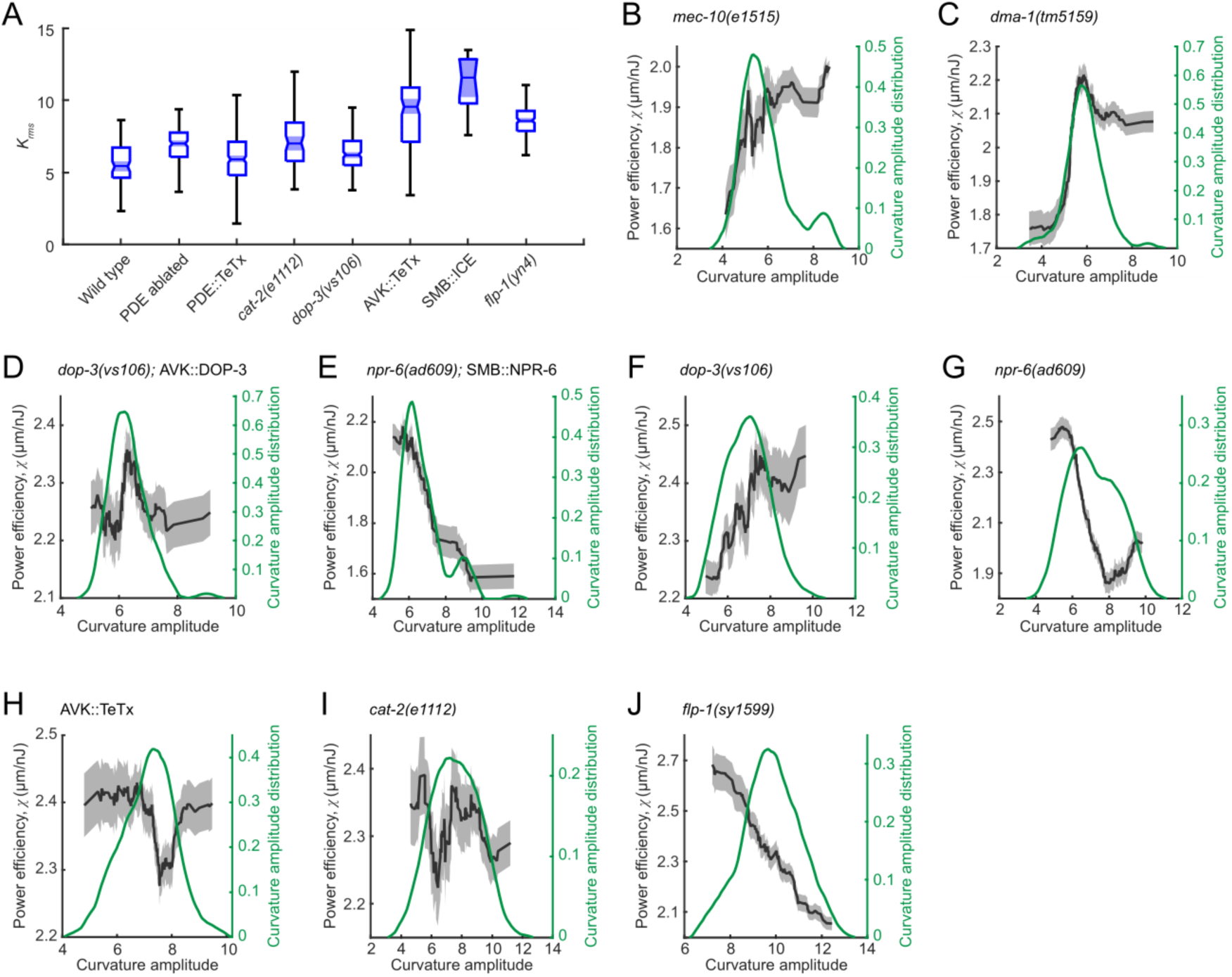
Biomechanical analyses of additional strains. (A) Locomotory amplitude in NGM solutions (1 mPa·s viscosity), measured from selective strains tested for CCR. Each box’s bottom, mid-line, and top are the data’s 25^th^, 50^th^, and 75^th^ percentiles. The whiskers go from the end of the interquartile range (between the bottom and top of the box) to the 1.5 times that range away from the bottom or top of the box. Data points beyond the whisker length represent outliers marked as red crosses. N = 80-150 forward moving bouts (10 s duration) from 10-13 animals per group. (B-J) Power efficiency as a function of curvature amplitude (black) and probability density function of curvature amplitude distribution (green) for strains with genotype *mec-10(e1515)* (A, n = 94 forward moving bouts from 16 animals), *dma-1(tm5159)* (B, n = 329 forward moving bouts from 20 animals), *dop-3(vs106);*AVK::DOP-3 (C, n = 200 forward moving bouts from 11 animals), *npr-6(ad609);*SMB::NPR-6 (D, n = 164 forward moving bouts from 10 animals), *dop-3(vs106)* (E, n = 150 forward moving bouts from 14 animals), *npr-6(ad609)* (F, n = 185 forward moving bouts from 11 animals), AVK::TeTx (G, n = 190 forward moving bouts from 12 animals), *cat-2(e1112)* (H, n = 138 forward moving bouts from 14 animals), and *flp-1(sy1599)* (I, n = 123 forward moving bouts from 9 animals). The 𝜒-𝑲 curve was obtained via a moving average along the x-axis with 1.5 in width, and the filled area represents a 95% confidence interval within the bin. Locomotion was assayed in dextran solutions with viscosity 120 mPa·s.

**Table S1.** List of worm strains.

| Strain ID | Source | Genotype |
| --- | --- | --- |
| BY602 | Blakeley lab | *dat-1(ok157); cat-2(e1112)* |
| BZ555 | CGC | *egls1[Pdat-1::GFP]* |
| CB1112 | CGC | *cat-2(e1112)* |
| CB1515 | CGC | *mec-10(e1515)* |
| CB156 | CGC | *unc-25(e156)* |
| CB1611 | CGC | *mec-4(e1611)* |
| CX4148 | CGC | *npr-1(ky13)* |
| DA702 | CGC | *eat-16(ad702)* |
| EJ1158 | CGC | *gon-2(q388)* |
| F41E7.3 | Gottschalk lab | *npr-6(tm1497)* |
| FG58 | CGC | *dop-4(tm1392)* |
| FQ2747 | Ringstad lab | *wzEx664[Pflp-1::TeTx; Pflp-1::mCherry]* |
| FQ2792 | Ringstad lab | *asic-1(ok415); del-3(ok2613); unc-8(wz211)* |
| FX5159 | Shen lab | *dma-1(tm5159)* |
| GS2477 | CGC | *arls37; cup-5(ar465); dpy-20(e1282)* |
| HBR507 | CGC | *flp-11(tm2706)* |
| JT603 | CGC | *gpb-2(sa603)* |
| JT734 | CGC | *goa-1(sa734)* |
| JT748 | CGC | *dgk-1(sy428)* |
| LX636 | CGC | *dop-1(vs101)* |
| LX702 | CGC | *dop-2(vs105)* |
| LX703 | CGC | *dop-3(vs106)* |
| LX704 | CGC | *dop-2(vs105); dop-3(vs106)* |
| LX705 | CGC | *dop-1(vs100); dop-3(vs106)* |
| LX706 | CGC | *dop-2(vs105); dop-1(vs100)* |
| LX734 | CGC | *dop-2(vs105); dop-1(vs100); dop-3(vs106)* |
| MT10549 | CGC | *tdc-1(n3421)* |
| MT1434 | CGC | *egl-30(n686)* |
| MT14984 | CGC | *tph-1(n4622)* |
| MT8504 | CGC | *egl-10(md176)* |
| NC279 | CGC | *del-1(ok150)* |
| NY16 | CGC | *flp-1(yn-4)* |
| NY2072 | CGC | *ynIs72[Pflp-1::GFP]* |
| PS8997 | Ringstad lab | *flp-1(sy1599)* |
| RM2221 | CGC | *egl-8(md1971)* |
| SWF141 | Flavell lab | *flvEx74[Pdat-1::CoChR; Pmyo-3::mCherry]* |
| SWF325 | Flavell lab | *flvEx133[Pdat-1::GtACR2-t2a-GFP; Pmyo-3::mCherry]* |
| SWF331 | Flavell lab | *flvEx127[Pdat-1::GCaMP6m; Pmyo-3::mCherry]* |
| TQ194 | CGC | *trp-2(sy691)* |
| TQ233 | CGC | *trpa-1(ok999)* |
| TQ296 | CGC | *trp-4(sy695)* |
| TU253 | CGC | *mec-4(u253)* |
| TV23560 | Shen lab | *otIs181[Pdat-1::mCherry; ttx-3::mCherry]; mals188[Pmir-288::GFP]* |
| VC1047 | CGC | *acd-3(ok1335)* |
| VC1210 | CGC | *del-8(ok1701)* |
| VM6365 | Villu Maricq lab | *akEx387[dat-1::GFP; dat-1::ICE]* |
| YX148 | This paper | *qhIs1[Pmyo-3::NpHR::eCFP]; qhIs4[Pacr-2::wCherry]* |
| YX287 | This paper | *cat-2(e1112); qhIs1[Pmyo-3::NpHR::eCFP]* |
| YX288 | This paper | *dop-3(vs106); qhIs1[Pmyo-3::NpHR::eCFP]* |
| YX289 | This paper | *cat-2(e1112); hpls199[Pmyo-3::ChR2::eGFP]* |
| YX290 | This paper | *dop-3(vs106); hpls199[Pmyo-3::ChR2::eGFP]* |
| YX291 | This paper | *dop-3(vs106); qhEx263[Pser-2-prom3::dop-3(+) + Punc-47::GFP]* |
| YX292 | This paper | *dop-3(vs106); qhEx264[Punc-47::dop-3(+) + Punc-47::GFP]* |
| YX293 | This paper | *dop-3(vs106); qhEx265[Pacr-2::dop-3(+) + Punc-47::GFP]* |
| YX294 | This paper | *dop-3(vs106); qhEx266[Pmyo-3::dop-3(+) + Punc-47::GFP]* |
| YX295 | This paper | *dop-3(vs106); qhEx267[Pacr-5::dop-3(+) + Punc-47::GFP]* |
| YX296 | This paper | *akEx387[dat-1::GFP; dat-1::ICE]; otIs181[Pdat-1::mCherry; ttx-3::mCherry]* |
| YX297 | This paper | *kyEx6101[Pdat-1::TeTx::sl2GFP]* |
| YX298 | This paper | *unc-54(e1092); flvEx127[Pdat-1::GCaMP6m; Pmyo-3::mCherry]* |
| ZM2509 | Zhen lab | *unc-9(fc16); hpEx803[Prgef-1::unc-9cDNA + Podr-1::GFP]* |
| ZM5398 | Zhen lab | *hpls199[Pmyo-3::ChR2::eGFP]* |
| ZX1291 | Gottschalk lab | *flp-1(yn4); zxEx474[Pflp-1(trc)::FLP-1::SL2::GFP]* |
| ZX2037 | Gottschalk lab | *npr-6(tm1497); zxEx850[Pflp-12::LoxP::LacZ::STOP::LoxP::NPR-6::SL2::GFP; Podr-2(18)::Cre]* |
| ZX2201 | Gottschalk lab | *dop-3(vs106); zxls20[Pdat-1::ChR2(H134R)::mCherry; Pmyo-2::mCherry];*<br>*zxEx1063[Pflp-1(trc)::DOP-3::SL2::GFP; Pmyo-3::CFP]* |
| ZX3058 | Gottschalk lab | *zxls29[Pflp-12::Cre; Podr-2(18)::LoxP::ICE; Pmyo-2::mCherry]* |

**Table S2.**
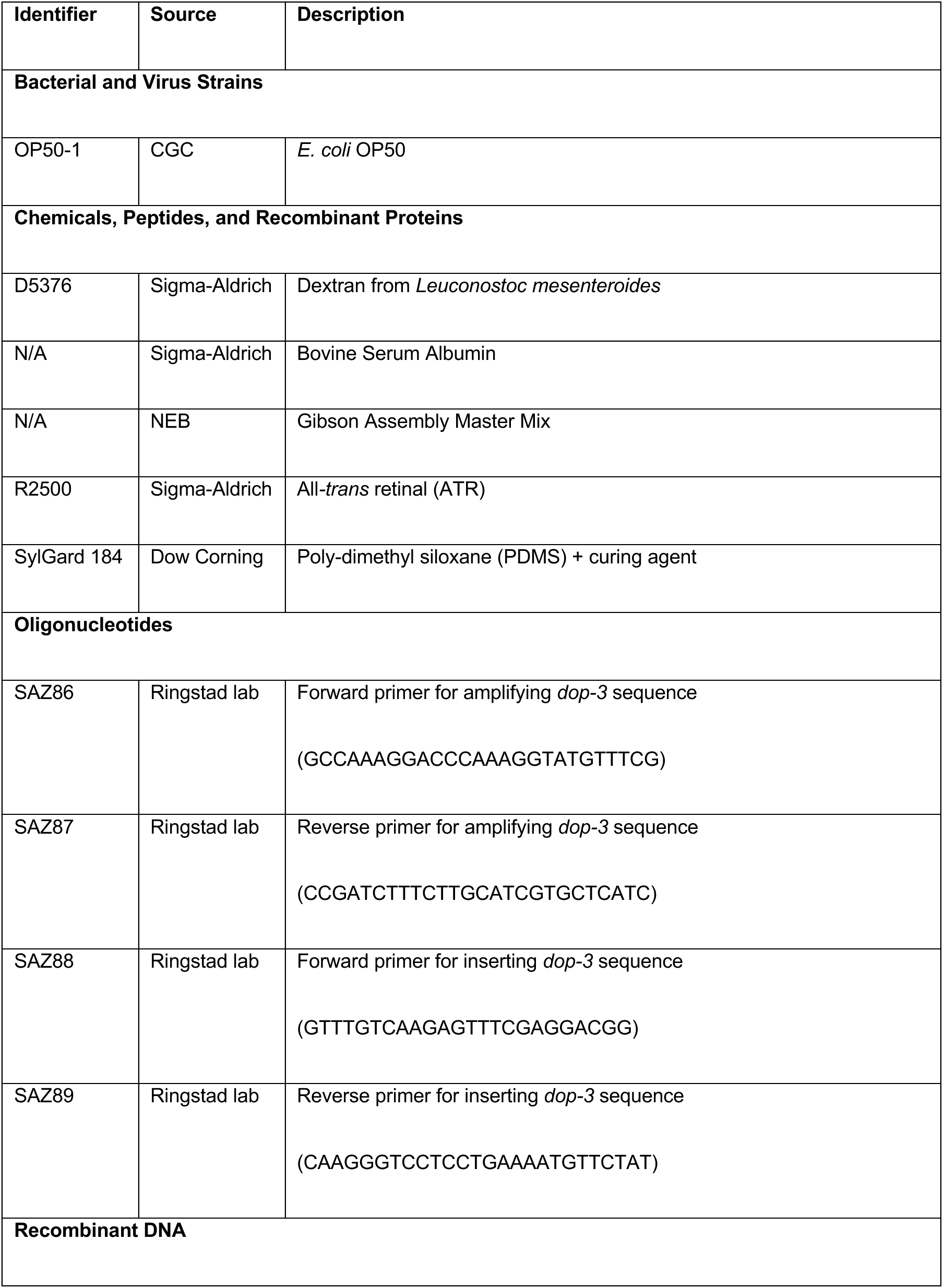
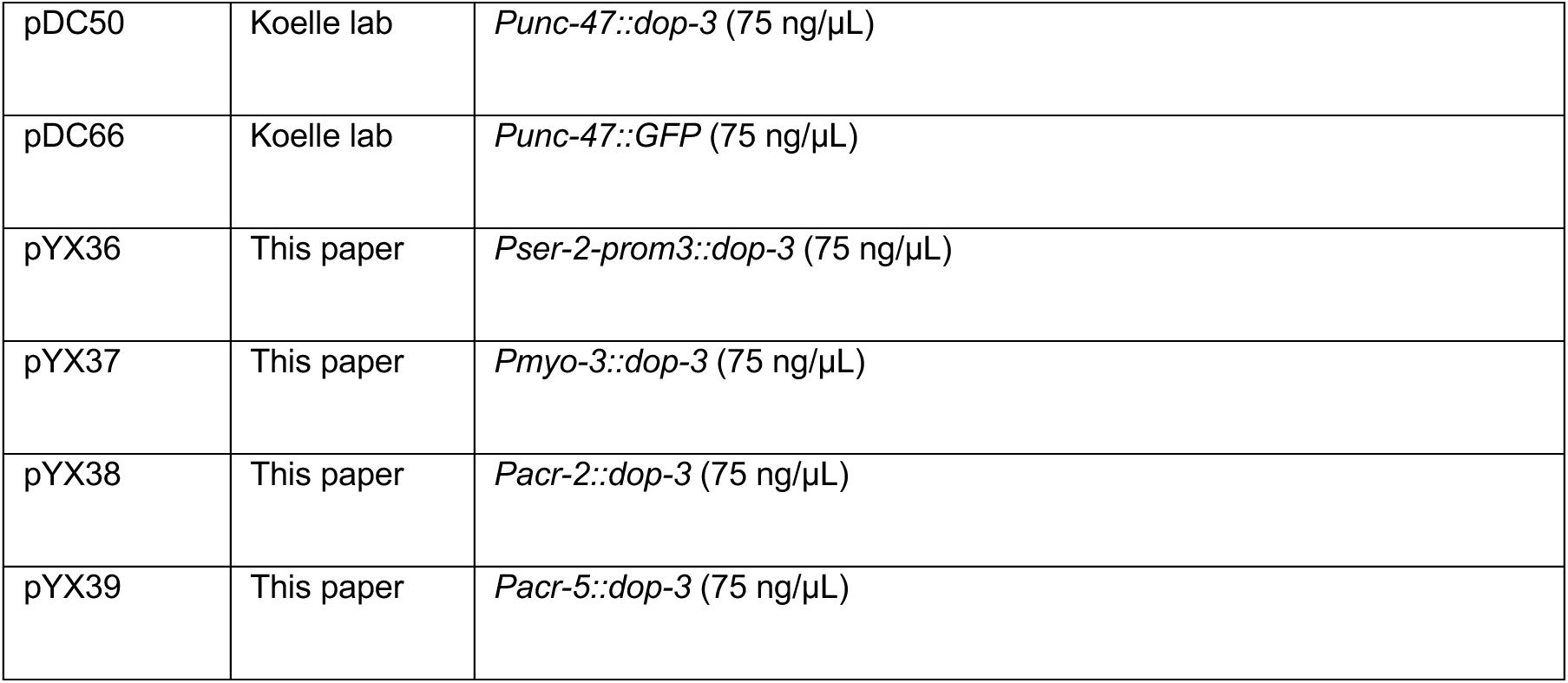
List of other reagents

**Movie S1 (separate file).** Transient optogenetic inhibition of muscles at various regions of the worm body.

**Movie S2 (separate file).** Transient optogenetic stimulation of muscle on one side of the middle region of the worm body.

**Movie S3 (separate file).** CCR of a wild-type animal.

**Movie S4 (separate file).** CCR of a *flp-1* mutant animal.

**Movie S5 (separate file).** CCR of animals with AVK ablated or AVK expressing tetanus toxin.

**Movie S6 (separate file).** CCR of a SMB::ICE transgenic worm.

## Notes

### Competing Interest Statement

The authors have declared no competing interest.

### Summary of Updates

We have made revisions throughout the document in response to reviewer comments and questions.

